# PPM1H phosphatase counteracts LRRK2 signaling by selectively dephosphorylating Rab proteins

**DOI:** 10.1101/711176

**Authors:** Kerryn Berndsen, Pawel Lis, Wondwossen Yeshaw, Paulina S. Wawro, Raja S. Nirujogi, Melanie Wightman, Thomas Macartney, Mark Dorward, Axel Knebel, Francesca Tonelli, Suzanne R. Pfeffer, Dario R. Alessi

## Abstract

Mutations that activate LRRK2 protein kinase cause Parkinson’s disease. LRRK2 phosphorylates a subset of Rab GTPases within their Switch-II motif controlling interaction with effectors. An siRNA screen of all protein phosphatases revealed that a poorly studied protein phosphatase, PPM1H, counteracts LRRK2 signaling by specifically dephosphorylating Rab proteins. PPM1H knock out increased endogenous Rab phosphorylation and inhibited Rab dephosphorylation. Overexpression of PPM1H suppressed LRRK2-mediated Rab phosphorylation. PPM1H also efficiently and directly dephosphorylated Rab8A in biochemical studies. A “substrate-trapping” PPM1H mutant (Asp288Ala) binds with high affinity to endogenous, LRRK2-phosphorylated Rab proteins, thereby blocking dephosphorylation seen upon addition of LRRK2 inhibitors. PPM1H is localized to the Golgi and its knockdown suppresses primary cilia formation, similar to pathogenic LRRK2. Thus, PPM1H acts as a key modulator of LRRK2 signaling by controlling dephosphorylation of Rab proteins. PPM1H activity enhancers could offer a new therapeutic approach to prevent or treat Parkinson’s disease.

## Introduction

LRRK2 is a large multidomain protein enzyme consisting of a Roc-Cor family GTPase followed by a serine/threonine kinase domain [1, 2]. Missense mutations within the GTPase domain (N1437H, R1441G/C, Y1699C) and kinase domains (G2019S, I2020T) hyper-activate LRRK2 protein kinase and cause Parkinson’s disease (PD) [3-5]. Mutations in LRRK2 are one of the most common genetic causes of familial Parkinson’s comprising ∼5% of familial Parkinson’s, and ∼1% of sporadic Parkinson’s patients [6, 7]. In terms of clinical presentation and late age of onset, LRRK2 mediated Parkinson’s closely resembles the common sporadic form of the disease affecting the vast majority of patients [8]. The G2019S mutation, located within the magnesium binding motif of the kinase domain, is the most common LRRK2 mutation and activates LRRK2 kinase activity around 2-fold [9-11]. Penetrance of the G2019S and other pathogenic LRRK2 mutations is incomplete [12, 13]. There is emerging evidence that the LRRK2 pathway is hyperactivated in some patients with idiopathic PD [14]. Pharmaceutical companies have developed LRRK2 inhibitors for treatment and prevention of PD [5] and clinical trials have commenced and/or are planned (see https://clinicaltrials.gov/ NCT03976349 (BIIB094) and NCT03710707 (DNL201)).

Recent studies have revealed that LRRK2 phosphorylates a subgroup of Rab proteins (Rab3A/B/C/D, Rab8A/B, Rab10, Rab12, Rab29, Rab35, and Rab43) at a conserved Thr/Ser residue (Thr73 for Rab10), located at the center of the effector binding switch-II motif [15-17]. Consistent with Rab proteins comprising disease-relevant substrates, all established pathogenic mutations enhance LRRK2 mediated Rab protein phosphorylation in a manner that is blocked by diverse LRRK2 inhibitors [15, 16, 18]. LRRK2 phosphorylation of Rab proteins blocks the ability of Rab proteins to interact with cognate effectors such as GDI and guanine nucleotide exchange factors, thereby trapping the phosphorylated Rab protein in the GTP bound state on the membrane where it has been phosphorylated [15, 19]. Recent work identified a novel group of effectors including RILPL1 and RILPL2 that bind preferentially to LRRK2 phosphorylated Rab8 and Rab10 [16]. LRRK2 phosphorylated Rab8A and Rab10, in complex with RILPL1/RILPL2, inhibit the formation of primary cilia that are implicated in controlling a Sonic hedgehog-driven neuroprotective pathway that could provide a mechanism by which LRRK2 is linked to PD [20]. Other research has revealed that components implicated in PD including Rab29 [21, 22] and VPS35 [23] also regulate phosphorylation of Rab proteins via LRRK2.

The protein phosphatase(s) that act on LRRK2 phosphorylated Rab proteins have not been identified, but appear to be highly active, as treatment of cell lines with LRRK2 inhibitors induce rapid dephosphorylation of Rab10 within 1-2 min [18]. In contrast, dephosphorylation of LRRK2 at a well-studied potential autophosphorylation site (Ser935) occurs much more slowly in response to LRRK2 inhibitor administration, requiring up to 90 min [18]. Only a small proportion (around 1%) of the total cellular Rab proteins are phosphorylated [15, 18]. In this study, we set out to identify and characterize the protein phosphatase(s) that counteract LRRK2 signalling by dephosphorylating LRRK2-modified Rab proteins.

## Results

### PP1 or PP2A phosphatases do not dephosphorylate Rab proteins

We first explored whether PP1 (Protein Phosphatase-1) or PP2A (Protein Phosphatase-2A) previously implicated in regulating LRRK2 [24, 25], control Rab phosphorylation. Wild type and LRRK2[R1441C] knock-in mouse embryonic fibroblasts (MEF cells) were treated with 0.1 µM calyculin-A (PP1 inhibitor) or 1 µM Okadaic acid (PP2A inhibitor) [26], and neither of these agents had a significant impact on basal phosphorylation of Rab10 (Thr73) or Rab12 (Ser105) or dephosphorylation following administration of the well-characterized highly selective and potent LRRK2 inhibitor termed MLi-2 [27] (Figure 1). Consistent with PP1 phosphatase regulating dephosphorylation of LRRK2 at Ser935 [24], calyculin-A stimulated Ser935 phosphorylation, particularly in the LRRK2[R1441C] MEF cells. It also prevented dephosphorylation of Ser935 following administration of MLi-2 (Figure 1). At 1 µM, Okadaic acid is likely to partially inhibit PP1 [26]; this concentration also blocked MLi-2 from inducing dephosphorylation of Ser935. Calyculin-A, but not okadaic acid promoted phosphorylation of the PP1 substrate MYPT1 (Thr853) [28]. Calyculin-A and okadaic acid also activated the S6K1 kinase pathway as judged by increase in S6K1 phosphorylation (Thr389, mTORC1 site) that is regulated by PP1 and PP2A [29, 30] (Figure 1).

**Figure 1.**
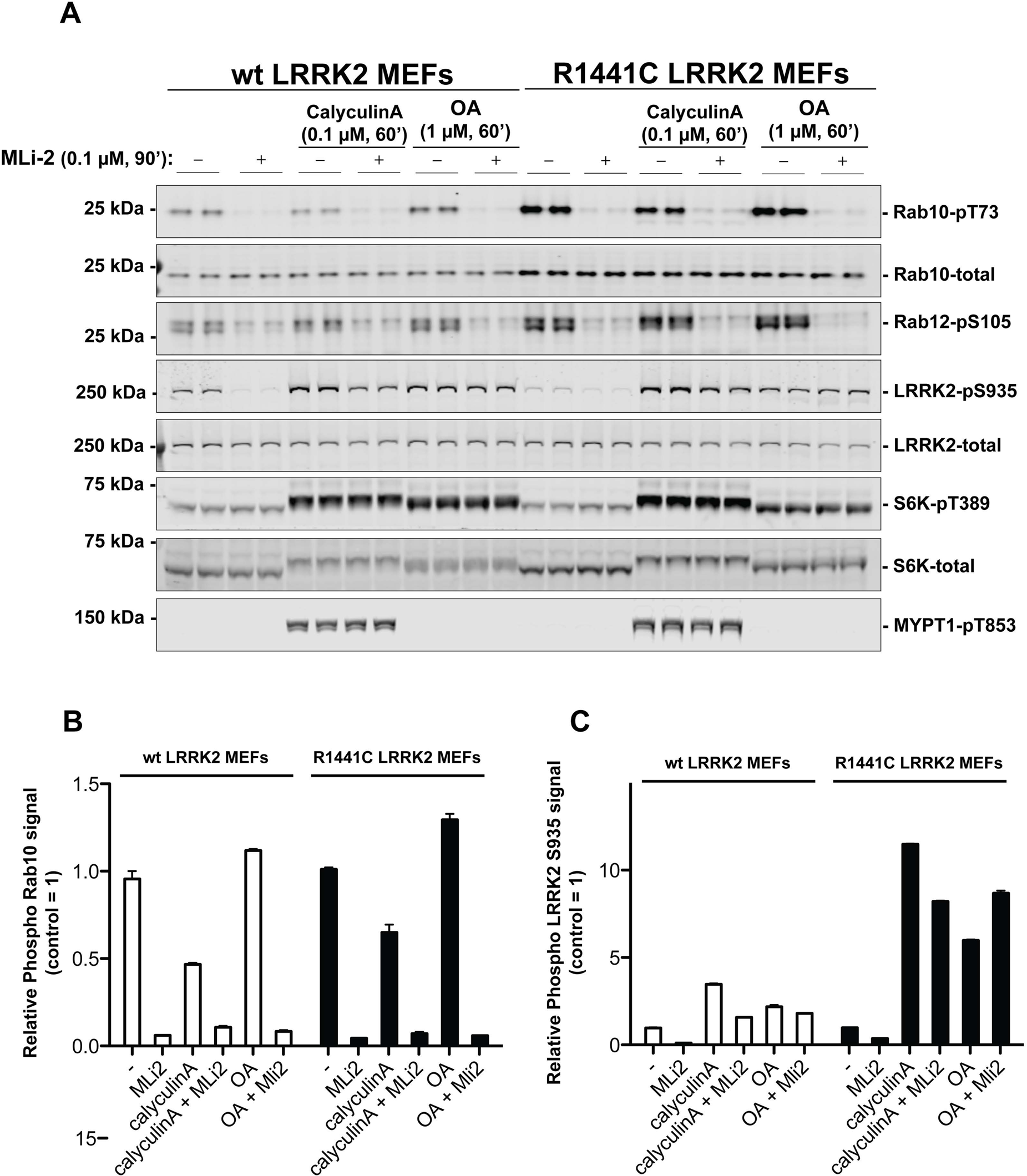
**PP1 or PP2A phosphatases do not dephosphorylate Rab proteins**. Littermate-matched wild-type and R1441C LRRK2 MEF cells were pre-treated with or without 100 nM MLi-2 for 90 min before administration with either 100 nM Calyculin A or 1 µM Okadaic Acid for further 60 minutes. Floating and adhering cells were lysed, and 10 µg of whole cell extract subjected to quantitative immunoblot analysis with the indicated antibodies (all at 1 µg/ml). (A) The membranes were developed using the LI-COR Odyssey CLx Western Blot imaging system. (B and C) Immunoblots were quantified for phospho-Thr73 Rab10/total Rab10 ratio (B) and phospho-Ser935 LRRK2/total LRRK2 ratio (C) using the Image Studio software. Data are presented relative to the phosphorylation ratio observed in cells treated with no MLi-2 and no phosphatase inhibitor, as mean ± SD.

### siRNA screen to identify phosphatases that regulate Rab10 phosphorylation

To identify protein phosphatases that act on LRRK2-phosphorylated Rab10, we undertook a phosphatase focused siRNA screen in human A549 adenocarcinomic alveolar basal epithelial cells that express endogenous LRRK2 [31]. We assembled a library consisting of 322 siRNAs (Dharmacon) targeting 264 phosphatase catalytic subunits (189 protein phosphatases, 26 halo acid dehalogenase phosphatases, 18 chloroperoxidases, 5 nucleoside-diphosphate-linked moiety-X phosphatases and 7 carbohydrate phosphatases) as well as 56 characterized regulatory subunits (Supplementary Excel File 1). Two replicate screens were undertaken in which cells were treated with siRNA targeting each phosphatase component for 72 h before cell lysis (Screen 1 & 2, Figures 2A & B). A third screen was also performed in which partial dephosphorylation of Rab10 was induced by adding MLi-2 inhibitor for 5 min prior to cell lysis (Screen 3, Figure 2A & C). Phospho-Thr73 Rab10 and total Rab10 were quantified in parallel employing a multiplex immunoblot assay. The ratio of phospho-Thr73 Rab10 versus total Rab10 was used to calculate the degree of Rab10 phosphorylation in each cell lysate. Levels of total LRRK2 and Ser935 phosphorylated LRRK2 were also analyzed. The results are summarized in Figure 2B (Screen 1 & 2) and Figure 2C (Screen 3). The primary immunoblotting data for each lysate is presented in Figure 2—Figure Supplement 1 (Screen 1), Figure 2— Figure Supplement 2 (Screen 2) and Figure 2—Figure Supplement 3 (Screen 3).

**Figure 2.**
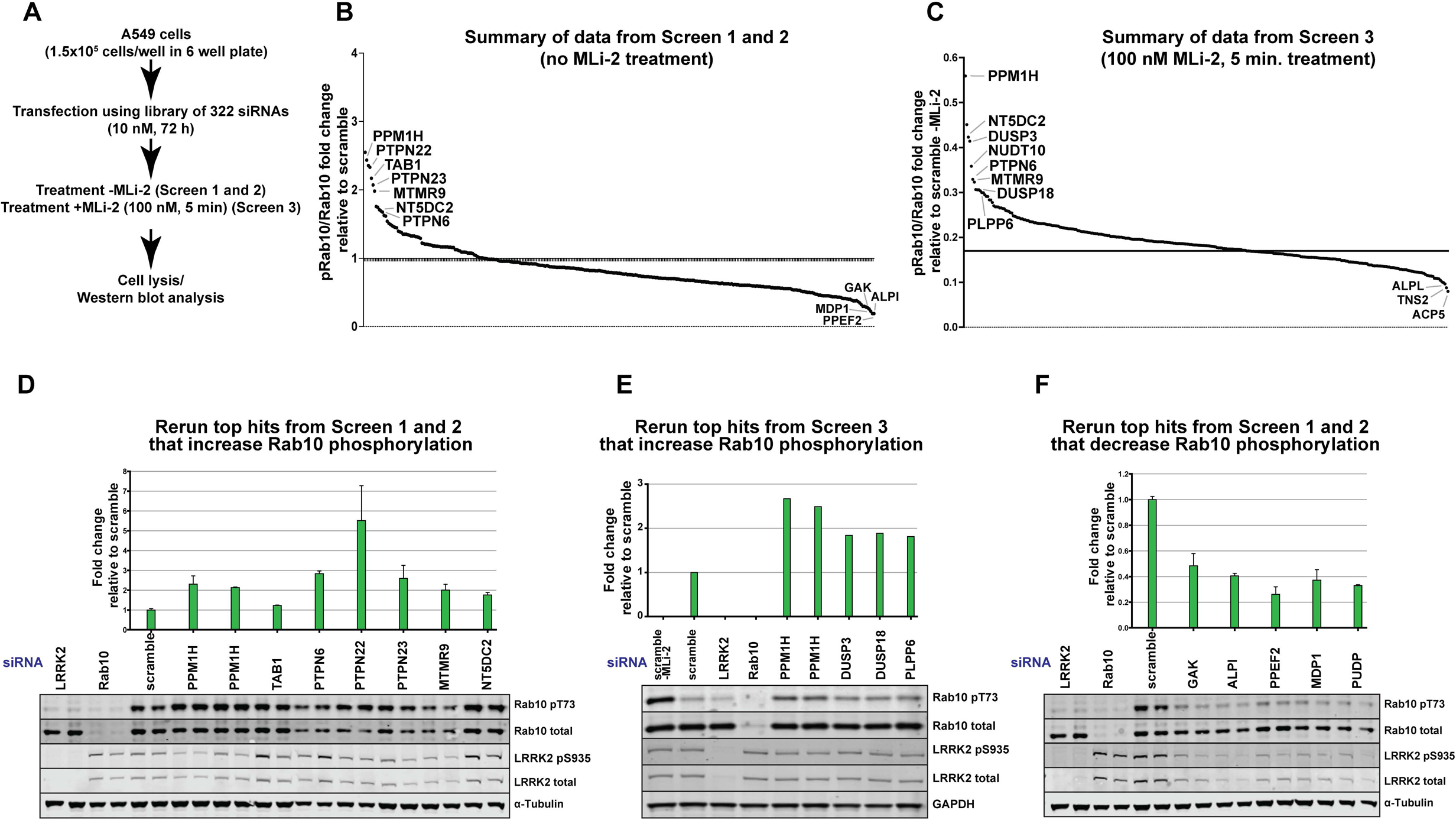
siRNA screens to identify phosphatases that regulate Rab10 phosphorylation. (A) Workflow of siRNA knockdown screens employed in this study. Human A549 cells were transfected with siRNA pools (Dharmacon) for 72h targeting 264 phosphatase catalytic subunits (189 protein phosphatases, 26 halo acid dehalogenase phosphatases, 18 chloroperoxidases, 5 nucleoside-diphosphate-linked moiety-X phosphatases and 7 carbohydrate phosphatases) as well as 56 characterized regulatory subunits. The list of phosphatase components targeted, and oligonucleotide sequences utilized is provided in Supplementary Excel File 1. Cells were lysed and immunoblotted for total LRRK2, LRRK2 pS935, total Rab10, and Rab10 pT73 and immunoblots developed using the LI-COR Odyssey CLx Western Blot imaging system. The ratio of phospho-Thr73 Rab10/total Rab10 in each sample was quantified using the Image Studio software. (B) Summary of results from Screen 1 and 2 (without MLi-2 pretreatment). Calculated ratio of phosphorylated Rab10 and total Rab10 relative the scramble control, ranked from highest increase in Rab10 phosphorylation to the strongest decrease (mean of the two replicates). (C) As in (B), summary of results from Screen 3 (in which cells were treated for 5 min with 100 nM MLi-2 treatment prior to cell lysis). (D) Human A549 cells were transfected with siRNA for 72 h with the indicated top hits from screen 1 and 2 that increased Rab10 phosphorylation. Cells were then lysed and immunoblotted with the indicated antibodies, analyzed as described above. Data are presented relative to the ratio of Rab10 phosphorylation/total Rab10 observed in cells treated with scrambled siRNA as mean ± SD. (E) As in (D) except the key hits that increased Rab10 phosphorylation in Screen 3 were reanalyzed. Cells were also treated with 100 nM MLi-2 prior to cell lysis. F) As in (D) except the key hits that decreased Rab10 phosphorylation in Screen 1 to 3 were reanalyzed.

**Figure 3.**
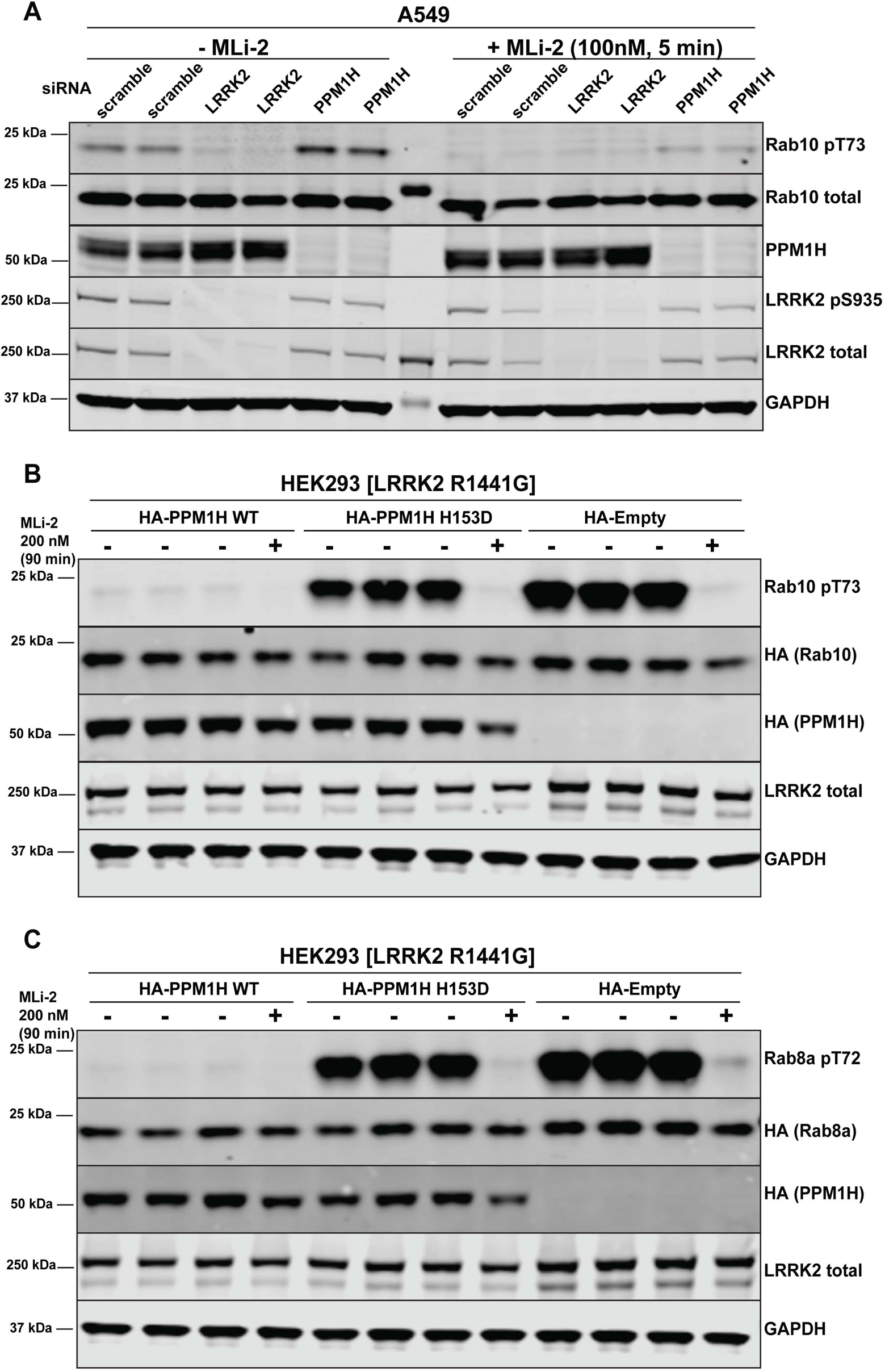
Further analysis of siRNA screen phosphatase hits. (A) Human A549 cells were transfected with the indicated siRNA pools (Dharmacon) for 72h. Cells were treated as indicated ± 100 nM MLi-2 for 5 min prior to lysis and 10 µg of whole cell extract subjected to quantitative LI-COR immunoblot analysis with the indicated antibodies (all at 1 µg/ml). (B & C) HEK293 cells were transiently transfected with constructs expressing the indicated components. 24 h post-transfection, cells were treated with ± 200 nM MLi-2 for 90 min and then lysed. 10 µg whole cell lysate was subjected to immunoblot analysis with the indicated antibodies at 1 µg/mL concentration and membranes analyzed as in (A).

The top hits that enhanced Rab10 protein phosphorylation over 1.5-fold in Screen 1 and 2 were PPM1H (Metal-dependent protein phosphatases [PPM] family serine/threonine phosphatase), TAB1 (PPM family catalytically inactive pseudophosphatase), PTPN6 (tyrosine phosphatase) PTPN22 (tyrosine phosphatase), PTPN23 (tyrosine phosphatase), MTMR9 (myotubularin lipid phosphatase) and NT5DC2 (nucleoside-diphosphate-linked moiety-X phosphatase) (Figure 2B). In screen 3 the tops hits were PPM1H, DUSP3 (dual specificity phosphatase), DUSP18 (dual specificity phosphatase) and PLPP6 (presqualene diphosphate phosphatase) (Figure 2C). We also identified 5 phosphatases (GAK, ALPI, PPEF2, MDP1, PUDP) that when knocked-down, significantly reduced Rab10 phosphorylation (Figure 2B).

### Further analysis of siRNA screen phosphatase hits

PPM1H was the only hit observed in all 3 screens and was also the top candidate in each of the screens. Human PPM1H possesses 514 residues encompassing a PPM type phosphatase domain with no other known functional motifs [32]. Repeat studies in multiple experiments confirmed that siRNA mediated depletion of PPM1H increased Rab10 phosphorylation, without affecting overall levels of Rab10 or LRRK2 (Figure 2D, 2E and 3A). Using a newly generated sheep polyclonal antibody, we established that the PPM1H expression was significantly lowered by the siRNA treatment (Figure 3A). Consistent with previous work [33], we observed that endogenous PPM1H, detected by immunoblot, migrates as a doublet of ∼55 kDa (reason for doublet unknown) with the top band migrating at the expected molecular weight for full length PPM1H. Expression of both forms of PPM1H was lowered by siRNA treatment. We also confirmed that siRNA knockdown of NT5DC2 (Figure 2D), DUSP3, DUSP18 and PLPP6 (Figure 2E) enhanced Rab10 phosphorylation without impacting Rab10, LRRK2 or LRRK2 Ser935 phosphorylation. However, knock-down of PTPN6, PTPN22, PTPN23 and MTMR9 significantly reduced Rab10 protein levels without affecting total LRRK2 (Figure 2D). siRNA depletion of TAB1 enhanced Rab10 protein phosphorylation to a marginal extent in repeat experiments (Figure 2D). Knock-down of GAK, ALPI, PPEF2, MDP1, and PUDP markedly decreased LRRK2 expression, likely accounting for the decreases in Rab10 phosphorylation observed (Figure 2F). These phosphatases were not investigated further in this study.

### Overexpression of PPM1H inhibits LRRK2-mediated Rab protein phosphorylation

Overexpression of wild type, but not catalytically inactive (His153Asp) PPM1H strikingly ablated the phosphorylation of Rab10 observed following overexpression of the LRRK2[R1441G] pathogenic mutant (Figure 3B). Similarly, overexpression of wild type, but not catalytically inactive PPM1H blocked phosphorylation of Rab8A (Figure 3C) as well as Rab3, Rab12, Rab35 and Rab43 (Figure 3— Figure Supplement 1A to 1D). In contrast, overexpression of PPM1H did not suppress phosphorylation of Rab8A at the Ser111 site induced following activation of the PINK1 kinase by addition of the mitochondrial uncoupler, CCCP [34] (Figure 3— Figure Supplement 1E). In addition, overexpression of PPM1H did not impact the phosphorylation of key markers of the Akt and AMPK signaling pathways (Figure 3— Figure Supplement 1F).

We also studied LRRK2-mediated phosphorylation of GFP-Rab10 stably expressed in wild type MEF cells, upon overexpression of wild type or catalytically inactive forms of PPM1H, by immunofluorescence microscopy (Figure 4). As observed previously, in cells not expressing either PPM1H form, phosphorylated Rab10 (detected using a phosphoRab10-specific antibody) was concentrated over peri-centriolar membranes, with a narrower localization than total Rab10 in these cells [20]. However, in cells expressing wild type PPM1H (but not its catalytically inactive forms), levels of phosphorylated Rab10 were markedly reduced if not abolished (Figure 4).

**Figure 4.**
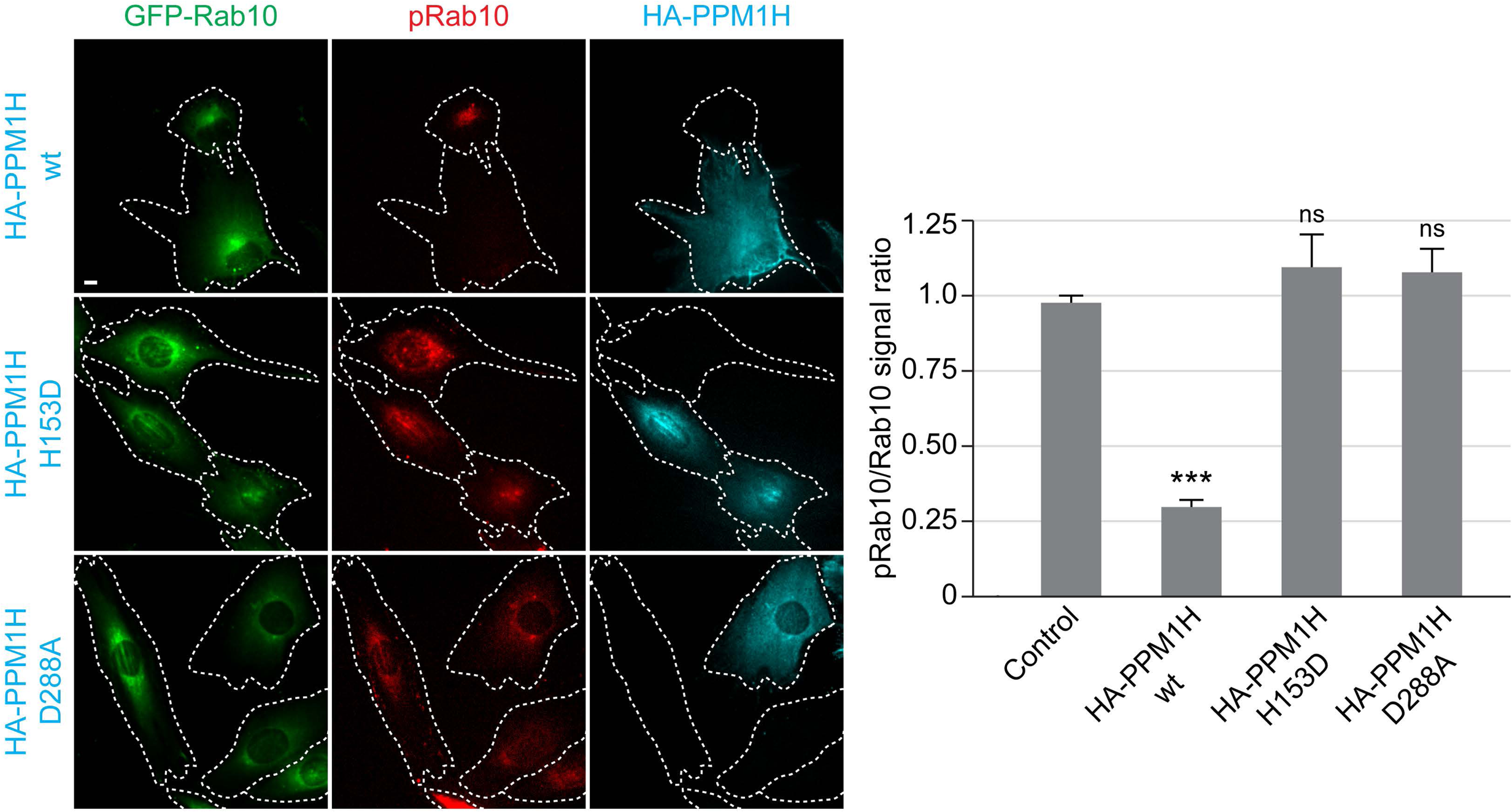
HA-PPM1H expression decreases pRab10 levels in MEF cells. Wild type MEF cells stably expressing GFP-Rab10 (green) were transiently transfected with plasmids encoding HA-PPM1H wild type, HA-PPM1H H153D, or HA-PPM1H D288A as indicated at left. After 24h the cells were subsequently fixed and stained with mouse anti-HA antibody (shown in blue) and rabbit anti-pRab10 antibody (shown in red). Scale bar,10 µm. Data were quantified by determining the ratio of pRab10 to GFP-Rab10 signal and normalizing these values to the ratio in cells not expressing PPM1H. For each condition, at least 20 cells were analyzed. ***, P < 0.001.

We next tested the two phosphatases most closely related to PPM1H termed PPM1J and PPM1M [32]. We found that PPM1M but not PPM1J suppressed Rab10 phosphorylation in cells albeit to a lower extent than PPM1H (Figure 5A). There are 20 PPM family phosphatases in humans (Figure 5B). In addition to PPM1M and PPM1J, we probed 10 other PPM members (TAB1, PPM1K, PPM1D, PPM1E, PPM1F, ILKAP, PPM1G, PPM1B, PPM1A, PHLPP1) and found that only PHLPP1, a membrane localized phosphatase [35], moderately reduced Rab10 phosphorylation (Figure 5C). Overexpression of other phosphatases identified in the siRNA screen had no (MTMR9, PTPN23) or only moderate effects (PLPP6, NT5DC2) on Rab10 phosphorylation (Figure 5— Figure Supplement 1).

**Figure 5.**
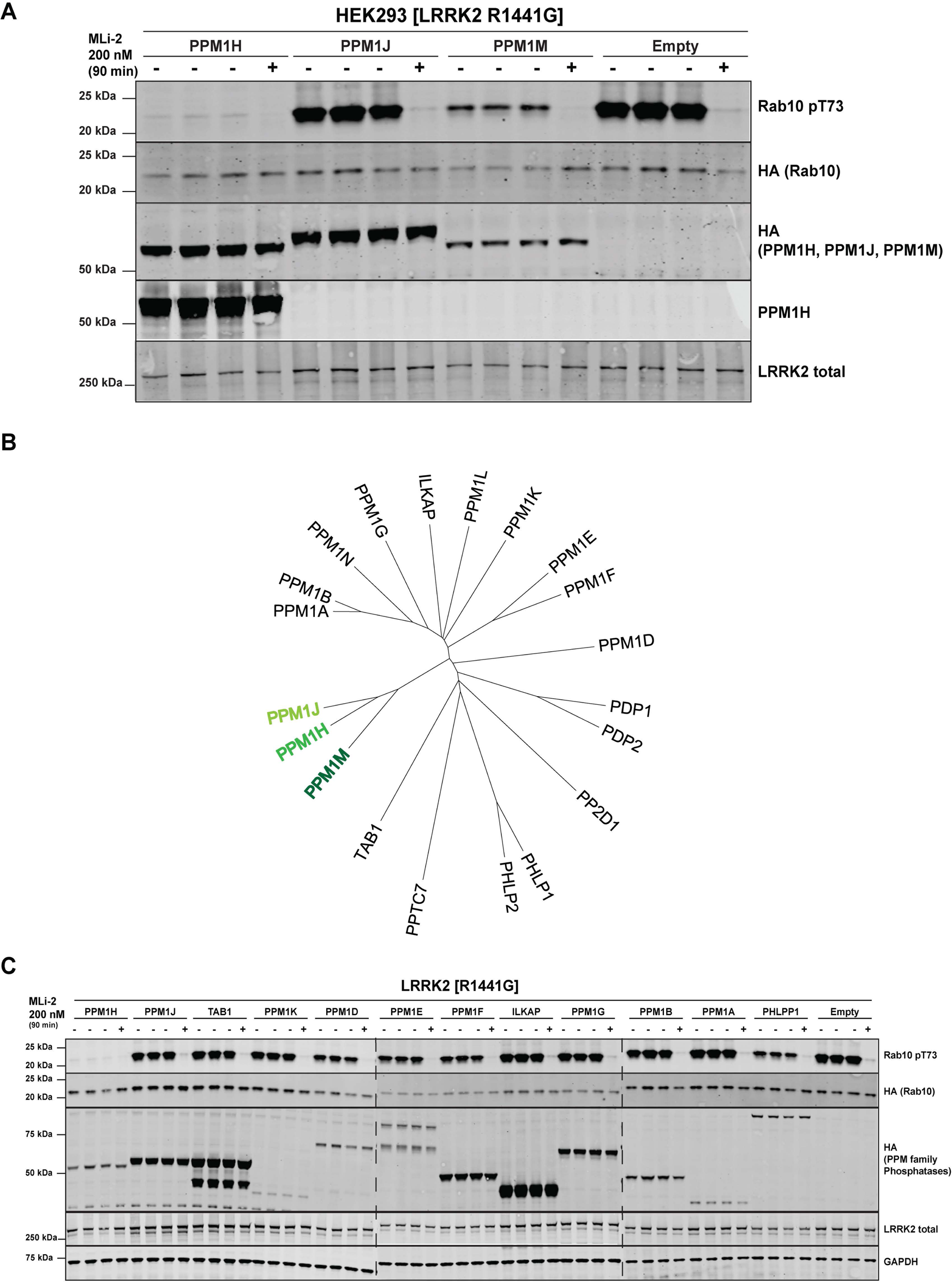
PPM1M, but not other PPM family members tested dephosphorylate Rab10. (A) HEK293 cells were transiently transfected with constructs expressing wild type PPM1H subfamily phosphatases (PPM1H, PPM1M and PPM1J). 24 h post-transfection, cells were treated with ± 200 nM MLi-2 for 90 min and then lysed. 10 µg whole cell lysate was subjected to immunoblot analysis with the indicated antibodies at 1 µg/mL concentration and membranes analyzed using the OdysseyClx Western Blot imaging system. Each lane represents cell extract obtained from a different dish of cells. (B) Phylogenetic tree of the 20 PPM family members generated using the MEGA7 software [66] . (C) As In (A) except that HEK293 cells were transiently transfected with constructs expressing the indicated PPM family members.

### CRISPR knock-out of full length PPM1H isoform enhances Rab protein phosphorylation

We next deployed a CRISPR/CAS9 gene editing approach to knock-out PPM1H in A549 cells. As mentioned above, immunoblotting studies suggested that there are potentially at least two forms of PPM1H expressed in cells. CRISPR-CAS9 constructs were designed that target exon-1 of the PPM1H gene and would be predicted to ablate expression of the full length PPM1H protein but may not impact expression of a shorter splice variant that lacks exon-1. We isolated 5 independent clones that demonstrated complete loss of the full length PPM1H protein by immunoblot analysis. These clones still possessed the lower species detected with the PPM1H antibody and indeed, expression of this form appeared to be upregulated in two of the cell lines (Figure 6A). Knock-out of the full length PPM1H species was sufficient to increase the basal level of phosphorylation of Rab10 in all 5-independent knock-out cell lines compared with two wild type cell clones that had been through all of the equivalent steps including clonal selection (Figure 6A). The rate of Rab10 dephosphorylation in PPM1H knock-out A549 cells following MLi-2 treatment was also markedly reduced compared with wild type cells (Figure 6B and 6C). In wild type cells, Rab10 was almost fully dephosphorylated within 1 minute of MLi-2 administration but remained fully phosphorylated for at least 2 minutes in PPM1H knock-out cells and remained more phosphorylated for up to 10 minutes. After 20 minutes, almost complete dephosphorylation of Rab10 was still observed in the knock-out cells, indicating that the full length PPM1H isoform that we have knocked out is not the only Rab10 protein phosphatase. In contrast, dephosphorylation of LRRK2 at Ser935 occurred at around 20 minutes after MLi-2 treatment and this was not affected by knock-out of the longer PPM1H isoform, consistent with distinct phosphatases acting to dephosphorylate LRRK2 and Rab10 (Figure 6B).

**Figure 6.**
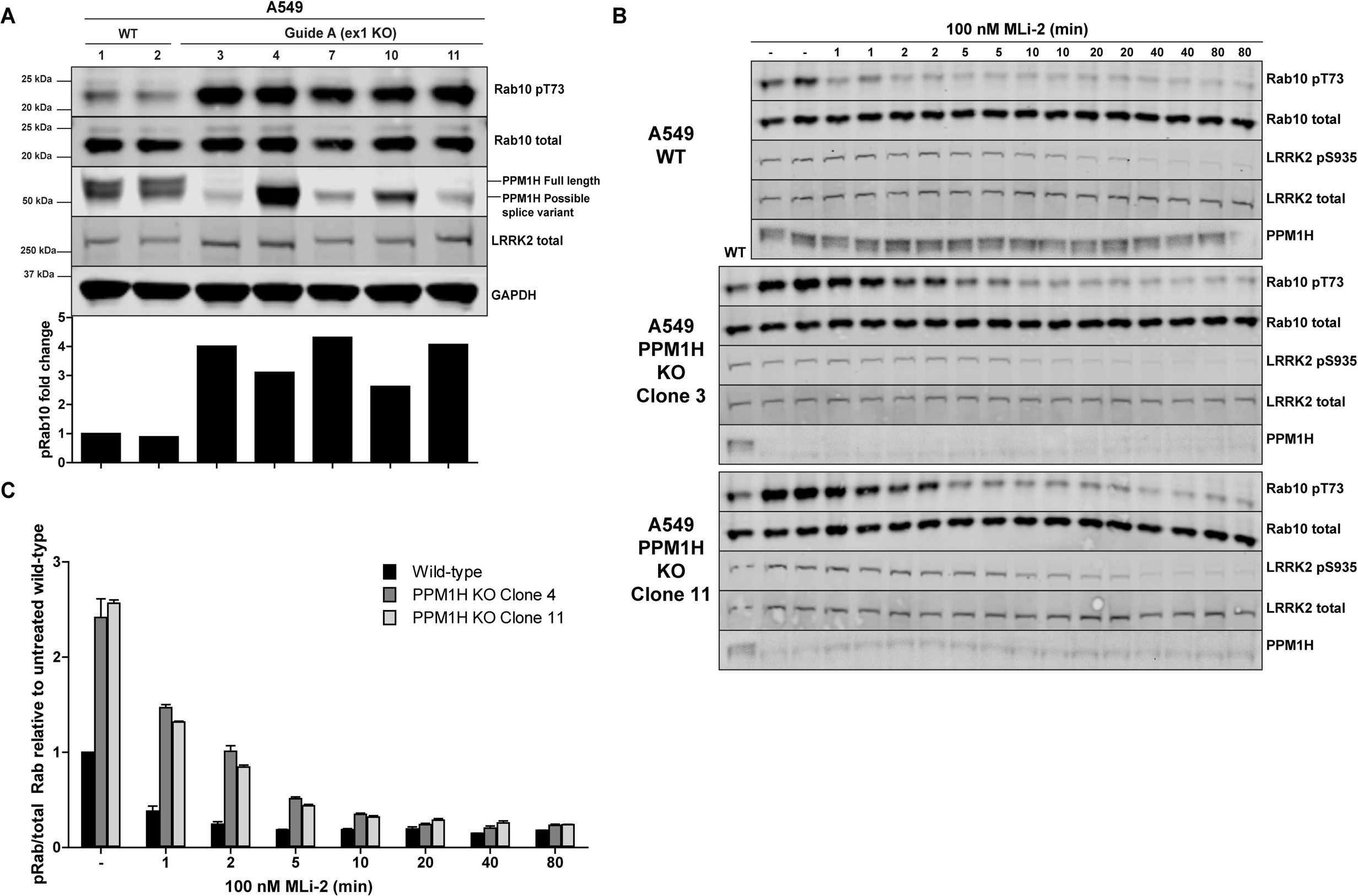
CRISPR knock-out of full length PPM1H isoform enhances Rab protein phosphorylation. (A) Two independent wild-type (WT 1 and WT 2) and five independent CRISPR/Cas9 PPM1H knock-out A549 cells (clone 3, 4, 7, 10, 11) that had all been through in parallel, single-cell sorting and expansion procedures were analyzed. 20 µg whole cell lysate was subjected to immunoblot analysis with the indicated antibodies at 1 µg/mL concentration and membranes analyzed using the OdysseyClx Western Blot imaging system. The ratio of phospho-Thr73 Rab10/total Rab10 was quantified using Image Studio software and data presented relative to the phosphorylation ratio observed in wild-type cells. (B) One Wild type and two independent CRISPR/Cas9 PPM1H knock-out A549 cells were treated ± 100nM MLi-2 inhibitor for the indicated times prior to cell lysis. 20 µg cell extract was analyzed and quantified (C) performed as in (A).

### PPM1H efficiently dephosphorylates Rab8A and Rab10 in vitro

To test whether PPM1H dephosphorylates Rab proteins in vitro, we produced recombinant RAB8A[Q67L] locked in the GTP-binding conformation, stoichiometrically phosphorylated at Thr72 (Figure 7—Figure Supplement 1 and 2). Phos-tag SDS-PAGE analysis was used to resolve phosphorylated and dephosphorylated Rab8A [18], providing a means to monitor direct dephosphorylation of Rab8A by recombinant PPM1H in biochemical experiments (Figure 7). Incubation of 2.5 µg phosphorylated Rab8A[Q67L] with PPM1H phosphatase for 30 min induced a dose dependent dephosphorylation with complete dephosphorylation of Rab8A[Q67L] observed with >40 ng of PPM1H and ∼50% dephosphorylation observed with 8 ng of PPM1H (Figure 7A). In parallel experiments, addition of 0.2 µg of catalytically inactive PPM1H[H153D] or catalytically inactive substrate trapping mutant PPM1H[D288A] (discussed below) markedly inhibited dephosphorylation of Rab8A[Q67L] (Figure 7A). PPM1H (40 ng) induced a time dependent dephosphorylation of Rab8A[Q67L]: 50% dephosphorylation was achieved within 10 minutes, and the reaction proceeded to completion by 80 minutes (Figure 7B). We also compared the rate that PPM1H dephosphorylated wild-type Rab8A complexed to GDP and GTPγS and found that both forms were similarly dephosphorylated (Figure 7A and 7B). In contrast, 0.2 µg PPM1M or PPM1J did not significantly dephosphorylate Rab8A[Q67L] under the same conditions in which 40 ng PPM1H quantitatively dephosphorylated Rab8A[Q67L] (Figure 7C and 7D).

**Figure 7.**
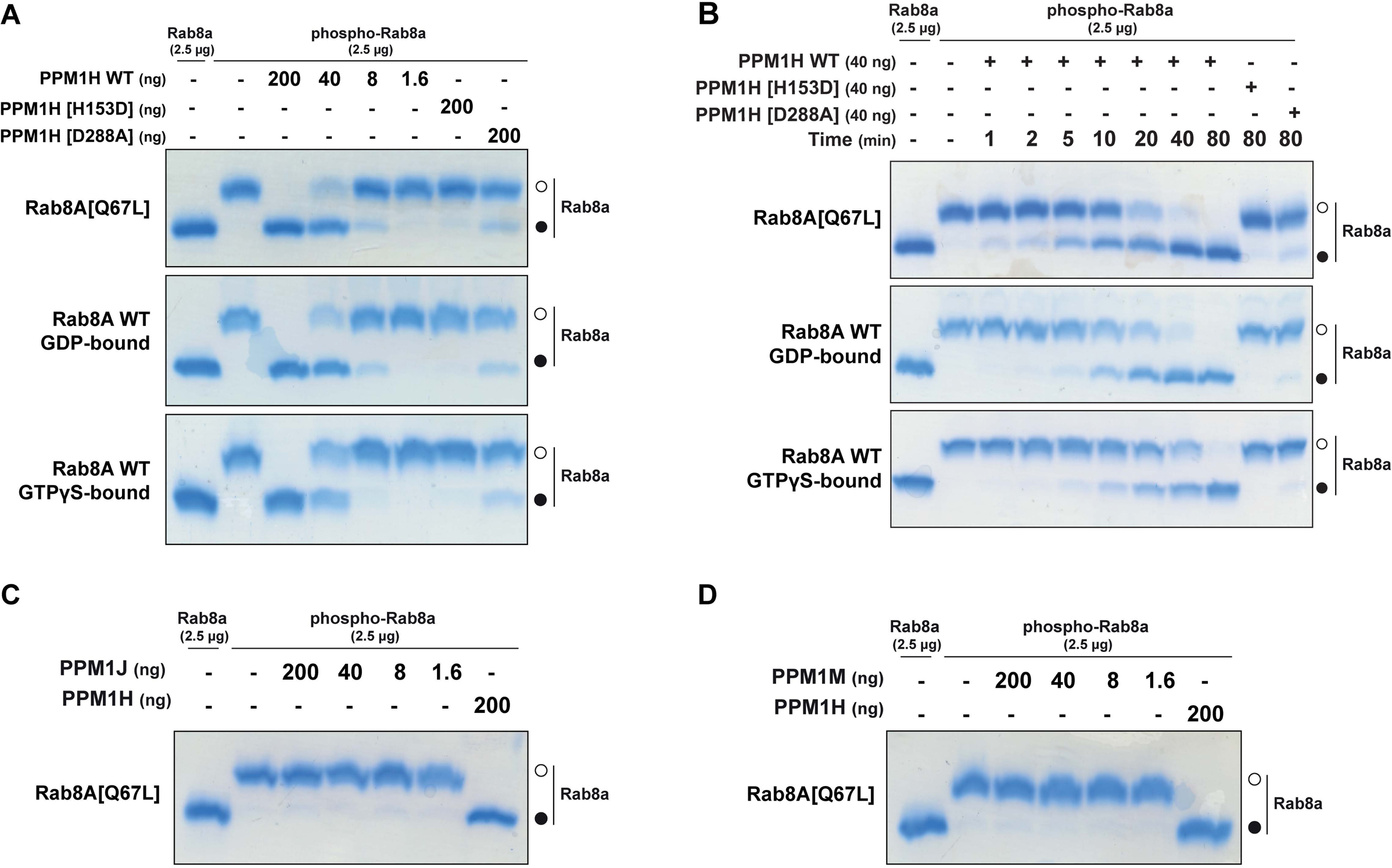
PPM1H dephosphorylates Rab8A[Q67L] in vitro. (A) The indicated amounts of recombinant wild type and mutant PPM1H (with a His-Sumo N-terminal tag, expressed in E.coli) were incubated in vitro with 2.5 µg pT72 phosphorylated Rab8A[1-181, Q67L-GTP bound conformation], 2.5 µg of Rab8A[1-181 wild-type, GDP-bound] or 2.5 µg of Rab8A[1-181 wild-type, GTP-bound] for 30 min in the presence of 10 mM MgCl2 in HEPES pH 7.0 buffer. Reactions were terminated by addition of SDS Sample Buffer and analyzed by phos-tag gel electrophoresis that separates phosphorylated and dephosphorylated Rab8A. Gel was stained with Instant Blue Coomassie. Bands corresponding to phosphorylated and non-phosphorylated Rab10 are marked with open (○) and closed (●) circles respectively. (B) As in (A) except that a time-course assay was performed using 2.5 µg pT72 phosphorylated Rab8A[1-181, Q67L-GTP bound conformation], 2.5 µg of Rab8A[1-181 wild-type, GDP-bound] or 2.5 µg of Rab8A[1-181 wild-type, GTP-bound] and 40 ng wild type or mutant PPM1H for the indicated times. (C) As in (A) except that PPM1J was assessed. (D) As in (A) except and PPM1M was assessed.

Analogous experiments were performed using wild type Rab10 locked in the GDP conformation that we were able to purify phosphorylated at Thr73 at a stoichiometry of about 60% (Figure 7-Figure Supplement 4 and 5). Dose dependent activity of PPM1H towards phosphorylated Rab10 followed a similar pattern to Rab8A, with complete dephosphorylation of 2.5 µg phospho-Rab10 observed after 30 minutes of incubation with >40 ng of PPM1H (Figure 7-Figure Supplement 3A). Time dependent activity assay of PPM1H towards Rab10 showed complete dephosphorylation after 80 minutes of incubation of 2.5 µg phospho-Rab10 with 40 ng of PPM1H (Figure 7-Figure Supplement 3B). 0.2 μg of purified PPM1J and PPM1M were also observed to weakly dephosphorylate Rab10 (Figure 7-Figure Supplement 3C and D).

### Identification of a substrate trapping PPM1H mutant

A recent study reported the crystal structure of a “substrate trapped” catalytically inactive mutant of PPM1A complexed with a phosphorylated peptide substrate [36]. This substrate trapping conformation was achieved by mutation of an Asp residue (Asp146 in PPM1A) that is conserved in 16 out of 20 PPM family phosphatases. We therefore explored whether mutation of the equivalent residue in PPM1H (Asp288) would also yield high affinity interaction with phosphorylated Rab proteins. We overexpressed wild type and PPM1H[D288A] in HEK293 cells expressing LRRK2[R1441G] and investigated whether endogenous phosphorylated Rab8A and Rab10 co-immunoprecipitate specifically with PPM1H[D288A]. Immunoblotting (Figure 8A) as well as high-resolution mass spectrometry (Figure 8C) analysis of the immunoprecipitates revealed that Rab8A and Rab10 were co-immunoprecipitated with the PPM1H[D288A] mutant, but not with wild type PPM1H. Phos-tag immunoblot analysis confirmed that it was the LRRK2 phosphorylated species of Rab8A and Rab10 that co-immunoprecipitated with PPM1[D288A] (Fig 8B).

**Figure 8.**
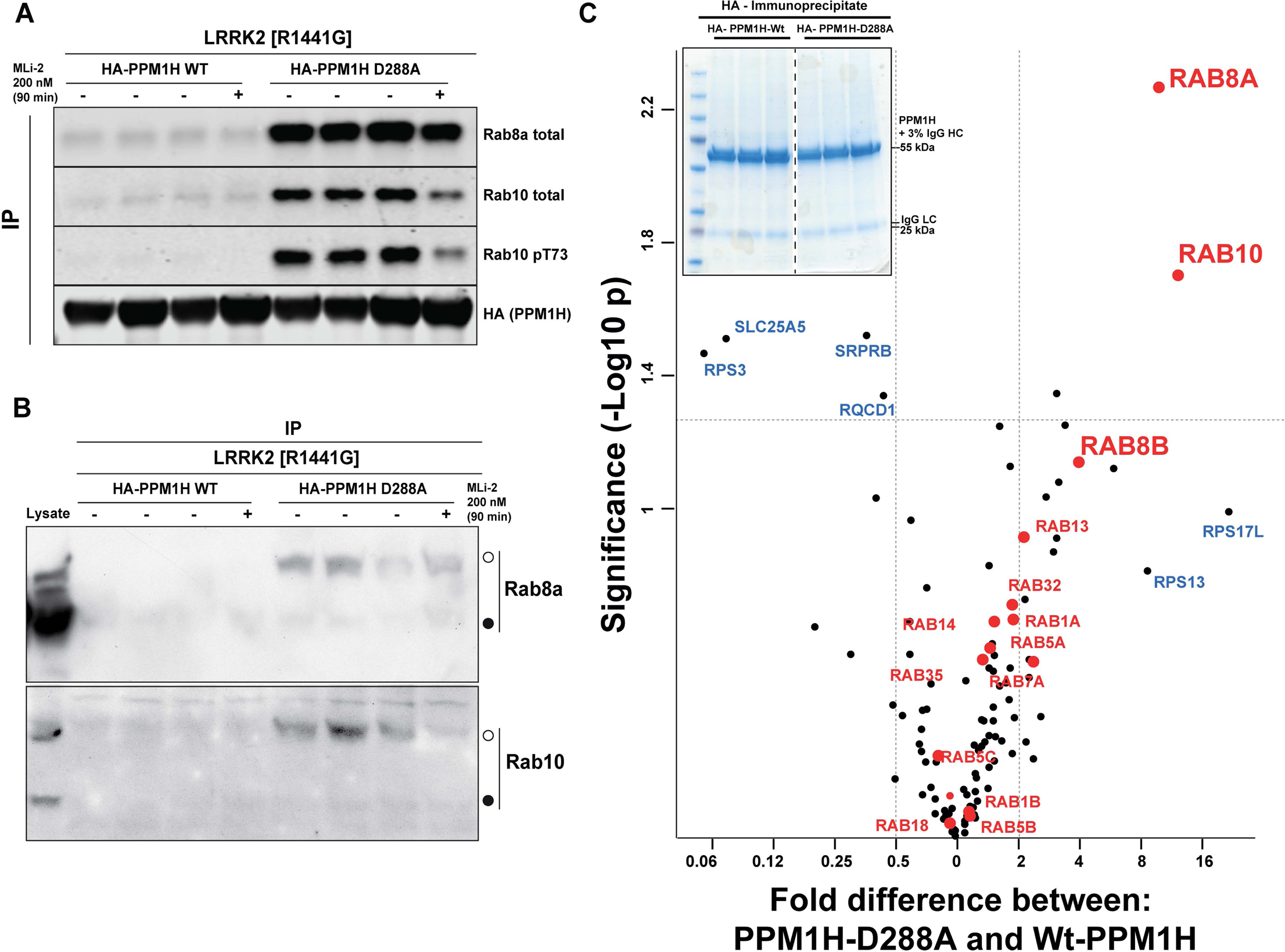
Identification of a substrate trapping mutant of PPM1H that forms a stable interaction with endogenous LRRK2 phosphorylated Rab8A and Rab10. (A & B) HEK293 cells were transiently transfected with constructs expressing Flag-LRRK2[R1441G] and either wild type HA-PPM1H or substrate trapping HA-PPM1H [D288A] mutant. 24 h post-transfection, cells were treated with ± 200 nM MLi-2 for 90 min lysed and subjected to a HA-immunoprecipitation and analyzed by either standard immunoblotting (A) or Phos-tag immunoblotting (B) with the indicated antibodies (1 µg/ml). Membranes were developed using Odyssey CLx Western Blot imaging. (C) As in (A) except that HA immunoprecipitates (n=3 for each condition) were subject to gel electrophoresis on a 4-12% Bis-Tris gel which is subsequently stained with Colloidal Coomassie blue USA (top left inset). The 20-30 kDa region of the gel that would encompass Rab proteins was excised, digested with trypsin and proteins present in each sample determined following quantitative mass spectrometry analysis. A Volcano plot analysis was performed to study differential enrichment of proteins that co-immunoprecipitate HA-PPM1H-D288A versus wild type PPM1H. Label free quantitation was undertaken, and protein intensities were subjected to Student-T test and permutation based false discovery rate of 5%. A score of >1.3 represented by the horizonal dotted line is considered significant. The vertical dotted lines represent samples whose abundance is 2-fold different in HA-PPM1H-D288A versus wild type PPM1H immunoprecipitates.

We also co-expressed PPM1H[D288A] with wild type, pathogenic or catalytically inactive LRRK2. This revealed that significantly higher levels of Rab10 co-immunoprecipitated with PPM1H[D288A] in cells expressing wild type or pathogenic LRRK2 compared with catalytically inactive LRRK2 (Figure 8 Figure Supplement 1A), consistent with PPM1H[D288A] interacting preferentially with LRRK2-phosphorylated Rab proteins. If PPM1H[D288A] forms a stable complex with phosphorylated Rab proteins in vivo, this would be expected to prevent their dephosphorylation following inhibition of LRRK2. Consistent with this, overexpression of PPM1H[D288A] and to a lesser extent PPM1H[D288E] suppressed dephosphorylation of Rab10 following treatment with 200 nM MLi-2 for 90 min, conditions that induce complete dephosphorylation of Rab proteins in the absence of PPM1H[D288A/E] overexpression (Figure 8— Figure Supplement 1B). Indeed, in cells expressing no phosphatase or the catalytically inactive PPM1H[H153D] mutant, Rab10 was almost completely dephosphorylated following 5-10 minutes of 200 nM MLi-2 treatment (Figure 8— Figure Supplement 2A and 2B). However, in cells expressing the PPM1H[D288A] substrate trapping mutant, dephosphorylation was markedly delayed and Rab10 was still significantly phosphorylated after 320 min 200 nM MLi-2 treatment (Figure 8— Figure Supplement 2C).

### PPM1H localizes to the Golgi

We used immunofluorescence microscopy of HA-PPM1H in RPE cells in relation to well characterized marker proteins including GCC185 (trans Golgi network), p115 and beta-COP (cis and medial Golgi), cation independent mannose 6-phosphate receptor (perinuclear late endosomes), and ACBD3 (medial Golgi; [37]; Figure 9). PPM1H was detected in the cytoplasm and also showed strong co-localization with the medial Golgi protein, ACBD3, as well as cis and medial Golgi markers, as quantified by Pearson’s correlation coefficient (Figure 9, left); this co-localization was retained upon fragmentation of the Golgi by nocodazole-induced microtubule depolymerization, the gold standard for membrane compartment localization (Figure 9, right). Thus, PPM1H is present in cytosol and also associates with the cis and medial Golgi complex.

**Figure 9.**
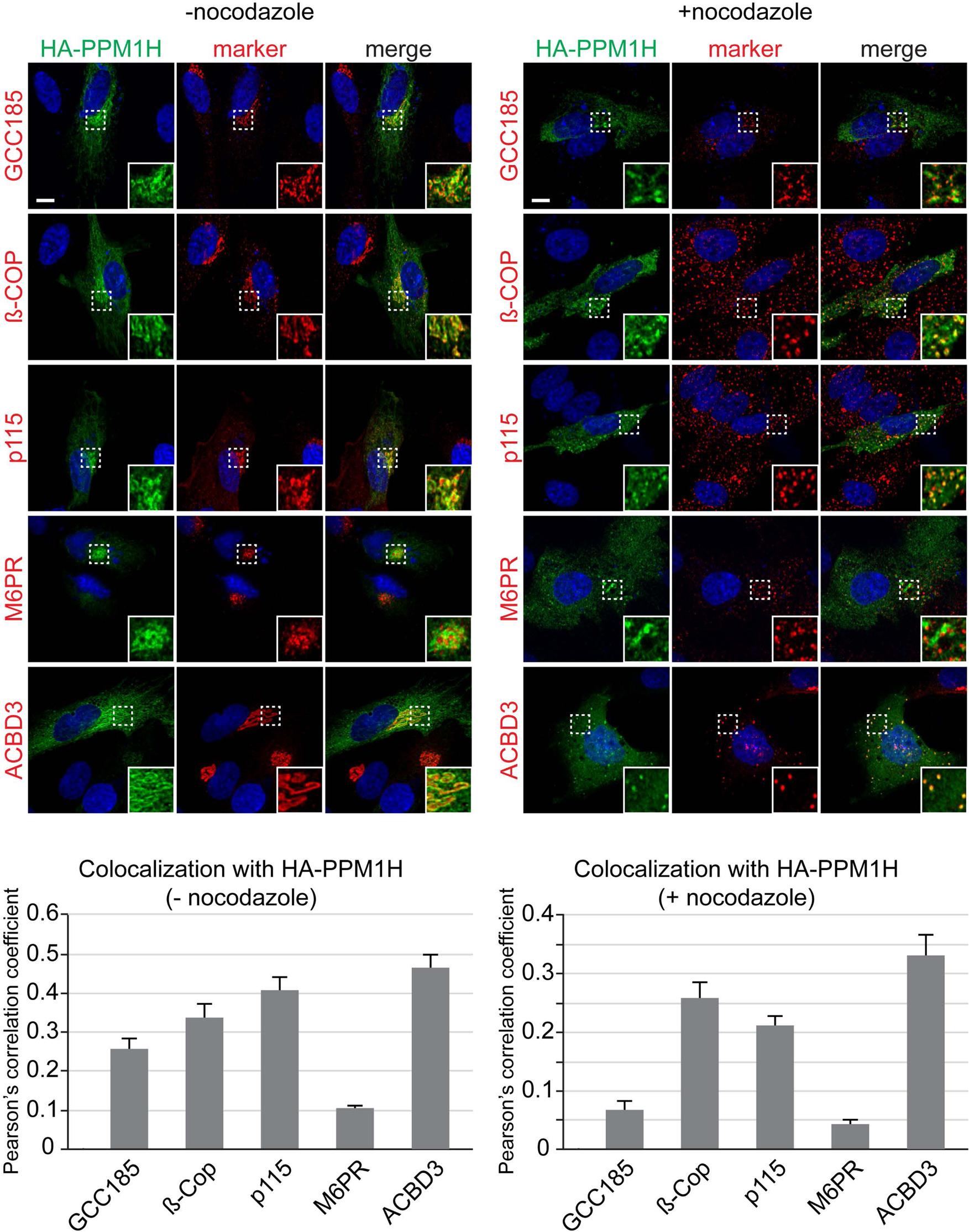
Localization of PPM1H in RPE cells. RPE cells were transiently transfected with HA-PPM1H plasmid. After 24h, cells were treated ± 20 µM nocodazole for 2 h before fixation. Cells were subsequently stained with mouse or rabbit anti-HA antibody (green) and the following Golgi markers (shown in red): rabbit anti-GCC185, rabbit anti-ß-COP, mouse anti-p115, mouse anti-cation independent mannose 6-phosphate receptor, and rabbit anti-ACBD3. Nuclei were stained with DAPI (blue). Scale bar, 10 µm. Shown are maximum intensity projections. For each condition at least 20 cells were analyzed.

### PPM1H knockdown suppresses primary cilia formation to the same extent as LRRK2 pathogenic mutations

We have shown previously that pathogenic LRRK2 suppresses cilia formation in cell culture and mouse brain [16, 20] in a process that requires Rab10 and RILPL1 proteins [20]. If Rab GTPase phosphorylation contributes to the regulation of cilia formation in wild type cells, depletion of PPM1H should also increase Rab phosphorylation, thereby inhibiting cilia formation. To test this, MEF cells were infected with lentiviruses encoding shRNAs to deplete PPM1H protein. As shown in Figure 10, depletion of PPM1H decreased cilia formation compared with control (scrambled shRNA sequence) infected cells, by a process that required LRRK2 as it was not seen when cells were cultured in the presence of the MLi-2 LRRK2 inhibitor. These experiments show that endogenous PPM1H protein contributes to the regulation of cilia formation in MEF cells and confirms a role for wild type LRRK2 protein in this important cellular process.

**Figure 10.**
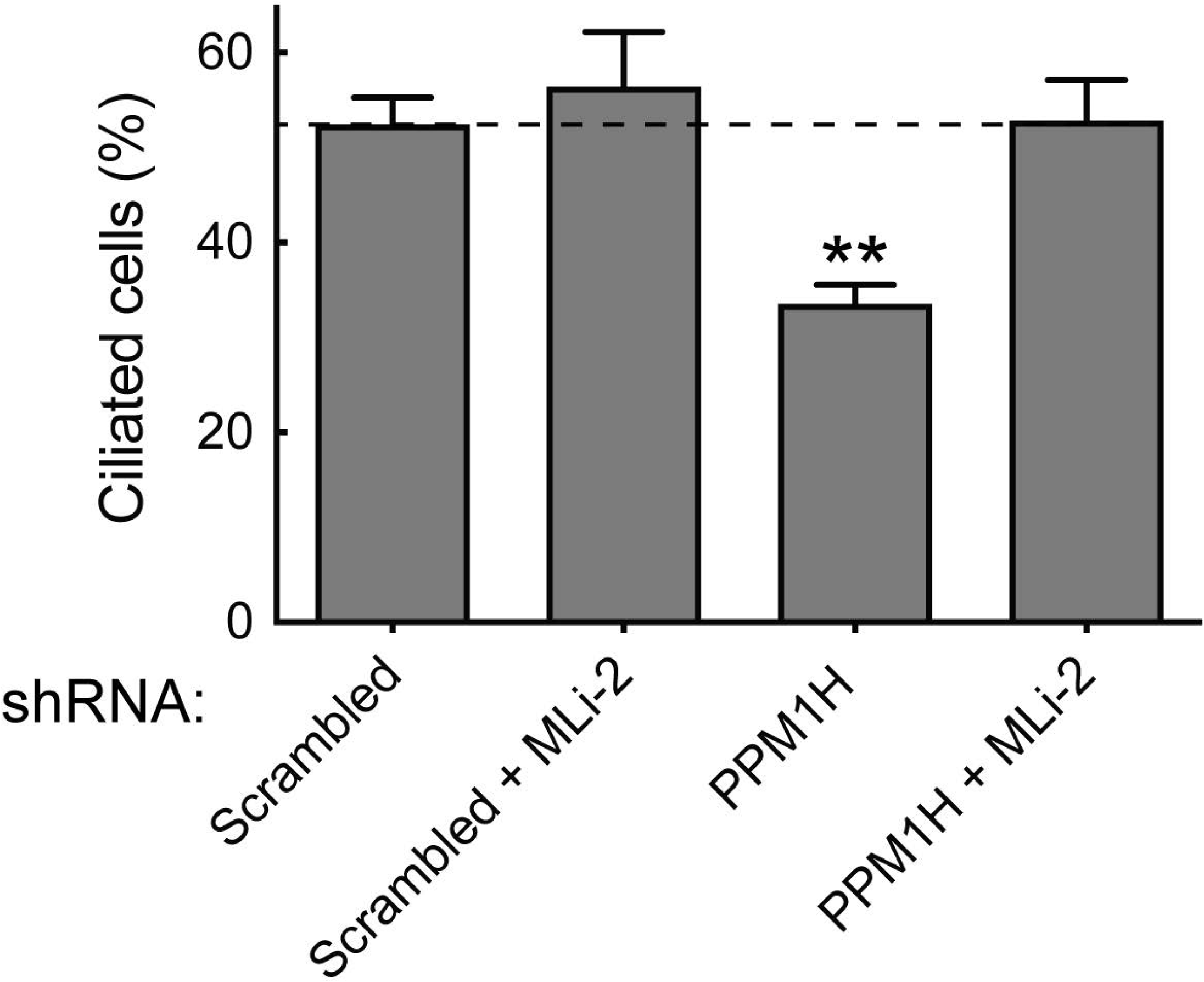
PPM1H depletion in wild type MEF cells decreases ciliation, consistent with increased Rab phosphorylation. Cells expressing the indicated shRNAs were assayed for ciliation 24h after serum starvation. Shown is the average of duplicate determinations made at identical cell confluency; error bars represent SEM. The experiment was replicated 4 times by two lab members; p=0.0018 ** (unpaired two-tailed t test, GraphPad PRISM software).

## Discussion

PPM phosphatases (previously known as PP2C phosphatases) are an ancient family of enzymes found in bacteria [38] and fungi [39]. Plants possess over 80 different PPM phosphatases that play important roles in hormone abscisic acid signaling and protection against stress [40, 41]. PPM phosphatases are evolutionarily unrelated to other serine/threonine protein phosphatases and there are 20 family members in humans (Figure 5B) [32]. These enzymes are characterized by their requirement for supplementation with millimolar concentrations of Mg2+ or Mn2+ ions for activity in vitro and are also insensitive to inhibition by okadaic acid or calyculin-A [42]. The structure of human PPM1A displays two tightly bound Mn2+ ions in the active site and a small subdomain, termed the Flap, located adjacent to the active site [43]. Crystal structures of bacterial [44] and plant [45] PPM homologs are nearly identical to that of the human PPM1A phosphatase core domain. In addition, a third, loosely associated magnesium ion, termed M3, appears to be required for optimal catalysis of all PPM family members [46].

PPM1H is a member of a group of 3 PPM phosphatases that have been termed the “PPM1H subfamily” [32]. This consists of PPM1H (URCC2, ARHCL1, NERPP-2C), PPM1J (PP2Cζ) and PPM1M (PP2Cη). PPM1H is most widely expressed in brain, but also found at varying levels in most other tissues including lung, kidney and spleen (Figure 11-Figure Supplement 1A to 1B). In blood cells, PPM1H is expressed at high levels in basophils and lower levels in neutrophils and is very low or not detected in most other cells including monocytes (Figure 11-Figure Supplement 2). PPM1M which is also able to dephosphorylate LRRK2-phosphorylated Rab10, was undetectable in most tissues and only observed at very low levels in the prostate gland and urinary bladder, despite mRNA levels being detected at similar levels to PPM1H in most tissues (Figure 11-Figure Supplement 1C to 1D). In the blood, PPM1M is most highly expressed in neutrophils (Figure 11-Figure Supplement 1C to 1D). Consistent with high expression in the brain, PPM1H was first discovered in a screen to identify protein phosphatases that are highly expressed in neuronal cell lines and in rat brains [47]. In future studies, it will be important to explore whether high levels of PPM1H in neuronal cells accounts for the relatively low levels of LRRK2 phosphorylated Rab10 that can be detected in brain in comparison with other tissues [31, 48].

The PPM1H subfamily phosphatases are conserved in animals from sponge to human [32]; invertebrates from sponge to ciona encode a single family member. The three copies found in mammals probably arose by two independent duplication events [32]. Human PPM1H possesses 514 residues encompassing a PPM type phosphatase domain with no other known functional motifs [32]. In previous work PPM1H was also reported to be overexpressed in colon adenocarcinoma and co-immunoprecipitated with the CSE1L apoptosis regulator that was suggested to comprise a substrate for PPM1H [49]. Another study found PPM1H confers trastuzumab resistance by promoting the dephosphorylation of p27 CDK inhibitor at threonine 187 [33]. It has also been suggested that patients whose tumors express low levels of PPM1H have reduced survival from colorectal cancer [50].

Our data reveal that PPM1H specifically dephosphorylates Rab proteins phosphorylated at their effector binding Switch region motif as it fails to dephosphorylate Rab8A protein phosphorylated at the distinct Ser111 site regulated by the PINK1 kinase also implicated in PD (Figure 3— Figure Supplement 1E) [34]. The specificity of PPM1H for phosphorylated Rab proteins is also highlighted by the substrate trapping PPM1H[D288A] mutant that binds to endogenous LRRK2 phosphorylated Rab8A, Rab8B and Rab10 with sufficient affinity to permit co-immunoprecipitation and suppress dephosphorylation (Figure 8). We also observed that PPM1H efficiently dephosphorylated Rab8A complexed to either GDP or GTPγS (Figure 7A and 7B), suggesting that it can recognize both these conformations. A priority for future work will be to elucidate the mechanism by which specificity of PPM1H for dephosphorylating LRRK2 phosphorylated Rab proteins is achieved. Structural analysis of the substrate-trapping mutant of PPM1H complexed to phosphorylated Rab8A will be highly informative. It will also be interesting to determine whether PPM1H possesses regulatory subunits that could control its activity, specificity and localization.

PPM1H is located both in cytosol and also on the cis and medial Golgi (Figure 9) and further work is needed to elucidate how PPM1H localizes to the Golgi and how it might translocate to other membranes and vesicles where LRRK2 phosphorylated Rab proteins reside. It is interesting to note that there is a significant pool of Rab10 on the Golgi and Rab8A on the Golgi and perinuclear compartments. Perhaps PPM1H’s Golgi localization serves to maintain those pools of Rab8A and Rab10 in functional forms to be able to perform functions other than cilia regulation. Another important goal will be to analyze the relationship between PPM1H localization and LRRK2 substrate selection in cells.

The PPM1H related, PPM1M enzyme (Figure 5B) also dephosphorylated Rab10 when overexpressed in cells, to a lower extent than PPM1H (Figure 5A). In corresponding biochemical studies, we also found that PPM1M displayed weak activity towards phosphorylated Rab10 (Figure 7-Figure Supplement 3D), but did not dephosphorylate Rab8A significantly (Figure 7D). In overexpression experiments PPM1J phosphatase displayed no activity towards dephosphorylating Rab10 (Figure 5A) although weak activity was observed in biochemical analysis (Figure 7-Figure Supplement 3C). Previous work revealed that at least 22 Rab proteins not regulated by LRRK2 are phosphorylated at their effector motifs [16], including Rab7A phosphorylated at Ser72 by TBK1 [51] or LRRK1 [52] and Rab1 that is phosphorylated at Thr75 by TAK1 [53]. It will be interesting to probe whether PPM1M, PPM1J or indeed PPM1H might dephosphorylate other Rab proteins. Out of the other 11 PPM family phosphatases tested, only PHLPP1 (PH domain leucine-rich repeat protein phosphatase), which possesses a membrane targeting PH domain, displayed weak ability to dephosphorylate Rab10 in an overexpression assay (Figure 5C).

As LRRK2 inhibitors still induce dephosphorylation of Rab10 in full length PPM1H knock-out A549 cells (albeit at a significantly reduced rate to the wild type cells) (Figure 6B), other protein phosphatase(s) must control Rab10 dephosphorylation in vivo. This could comprise PPM1M and/or other phosphatases. Our analysis also revealed that knock-down of a number of other phosphatases (TAB1, PTPN6, PTPN22, PTPN23, MTMR9, NT5DC2, DUSP3, DUSP18, PLPP6) modestly increased Rab10 protein phosphorylation (Figure 2D and 2E). These phosphatases include tyrosine phosphatases (PTPN6, PTPN22, PTPN23), dual specificity phosphatases (DUSP3, DUSP18), a lipid phosphatase (MTMR9), a catalytically inactive pseudophosphatase (TAB1), and metabolic phosphatases (NT5DC2 and PLPP6). None of these enzymes are known to dephosphorylate Ser/Thr residue phosphorylated proteins and it is possible that the observed actions of these phosphatases are indirect; further analysis will be required to probe any physiological significance. Interestingly, we also found that knock-down of a small group of phosphatases (GAK, ALPI, PPEF2, MDP1, PUDP) reduced LRRK2 levels leading to an inhibition of Rab10 protein phosphorylation (Figure 2F). One of these, GAK, is also known as Auxilin-2, and genome-wide association studies have revealed variants linked to PD to lie within or close by to the GAK gene [54]. It will be interesting to probe the mechanisms by which GAK as well as the other phosphatases impinge on the LRRK2 pathway.

In summary, our data provide compelling evidence that PPM1H acts to dephosphorylate Rab proteins in vivo and therefore counteracts LRRK2 signalling (Figure 11). This is supported by the findings that knock-down (Figure 3A) as well as knock-out (Figure 6) of PPM1H induce endogenous Rab10 protein phosphorylation and PPM1H directly dephosphorylates Thr72 phosphorylated Rab8A in vitro (Figure 7). Expression of PPM1H in cells abolishes phospho-Rab10 staining of wild type MEF cells (Figure 4). We identify a substrate trapping mutant of PPM1H that binds to endogenous LRRK2 phosphorylated-Rab8A as well as Rab10 and also protects Rab10 from becoming dephosphorylated following inhibition of LRRK2 (Figure 8). LRRK2 has been reported to regulate many biological responses such as autophagy [55], immune responses [56] as well as phagosome maturation [57], and a key question is whether LRRK2 controls these effects by phosphorylating Rab proteins or via an independent pathway. This issue of whether Rab proteins are involved in mediating the actions of LRRK2 can be addressed in the future by probing the impact of manipulating PPM1H levels. Indeed, we have used this approach to demonstrate that knock-down of PPM1H blocks primary cilia formation in a manner analogous to pathogenic, activating LRRK2 mutations (Figure 10). This is consistent with our previous work demonstrating that the ability of LRRK2 to inhibit primary cilia formation is indeed dependent on Rab8A and Rab10 protein phosphorylation [20]. In future work it will be important to explore whether PPM1H contributes to PD risk. For example, pathogenic mutants of LRRK2 are incompletely penetrant and a majority of patients may not develop PD [12, 13]. It would be critical to explore whether increased expression or activity of PPM enzymes protects patients with LRRK2 mutations from developing PD by enhancing Rab protein dephosphorylation. Conversely, reduced expression or activity of PPM1H phosphatase would be expected to promote Rab protein phosphorylation and enhance PD risk. Targeting PPM1H to increase its activity or expression in order to promote Rab protein dephosphorylation could be explored as a therapeutic strategy for preventing and/or treating LRRK2-mediated PD.

**Figure 11.**
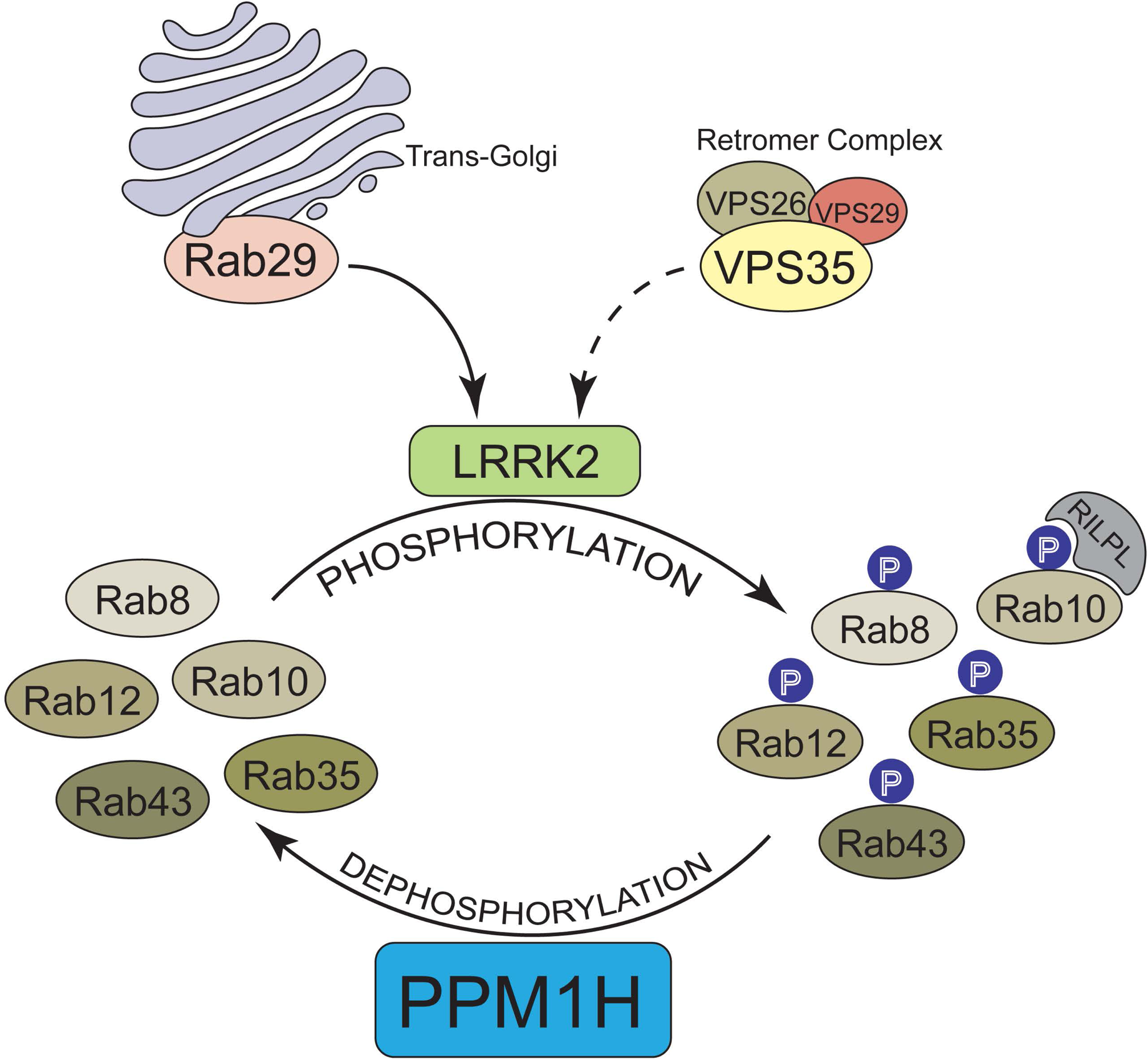
Model of how PPM1H counteracts LRRK2 signalling pathway by dephosphorylating Rab proteins.

## Material and Methods

### Cell culture and lysis

Wild-type and homozygous LRRK2 R1441C knock-in MEF cells isolated from littermate-matched mouse embryos at day E12.5 (described in [16]) were grown in DMEM containing 10% (by vol) fetal calf serum, 2 mM L-glutamine, 100 U/ml penicillin, and 100 mg/ml streptomycin supplemented with non-essential amino acids and 1 mM sodium pyruvate. A549, RPE and HEK293 cells were purchased from ATCC and cultured in DMEM containing 10% (by vol) fetal calf serum, 2 mM l-glutamine, 100 U/ml penicillin, and 100 mg/ml streptomycin. All cells were grown at 37oC, 5% (by vol) CO2 in a humidified atmosphere and regularly tested for mycoplasma contamination. Unless otherwise indicated cells were lysed in an ice-cold lysis buffer containing 50 mM Tris–HCl, pH 7.4, 1% (by vol) Triton X-100, 10% (by vol) Glycerol, 0.15 M NaCl, 1 mM sodium orthovanadate, 50 mM NaF, 10 mM 2-glycerophosphate, 5 mM sodium pyrophosphate, 1 µg/ml microcystin-LR, and complete EDTA-free protease inhibitor cocktail (Roche). Lysates were clarified by centrifugation at 20 800 g at 4°C for 10 min and supernatants were quantified by Bradford assay.

### Antibodies

To generate a polyclonal PPM1H antibody a sheep was immunized with full length PPM1H protein (His-SUMO tag cleaved off) followed by 3 further injections 28 days apart, with bleeds performed seven days after each injection. Antibodies were affinity-purified from serum using PPM1H protein. For immunoblotting analysis, the PPM1H sheep antibody was used at a final concentration of 1 μg/ml. We noted that incubation for 90 min at room temperature appeared to provide clearer results than overnight incubation at 4°C. The sheep PPM1H antibody is available from MRC PPU Reagents and Services (sheep number DA018 https://mrcppureagents.dundee.ac.uk/). Sheep polyclonal antibody detecting total Akt was purified at the University of Dundee (S695B). The Phospho-AMPKα (Thr172) (CST (40H9) #2535), AMPKα total (CST (D5A2) #5831) and Phospho-Akt Ser473 (CST #9271) were purchased from Cell Signaling Technology. Mouse anti-GAPDH total was from Santa Cruz Biotechnology (sc-32233). PINK1 total antibody was from Novus (BC100-494), OPA1 antibody was from BD (612607). Rabbit monoclonal antibodies for total LRRK2 (N-terminus) (UDD3) and phospho-Ser935 LRRK2 (UDD2) were expressed and purified at University of Dundee as described previously [58]. The recombinant MJFF-pRab10 (Thr73) and MJFF-pRab8 (Thr72) rabbit monoclonal antibodies were recently described [31] and are available from Abcam (#ab230261 and ab230260). The rabbit polyclonal Rab8A phospho-S111 antibody was obtained from Miratul Muqit (University of Dundee). Recombinant antibodies were used at 1 µg/ml final concentration for immunoblotting. The MJFF-total Rab10 mouse antibody was from nanoTools (0680-100/Rab10-605B11, www.nanotools.de) [31]and was used at 1 µg/ml final concentration. The phospho-Rab12 (S106) rabbit monoclonal antibody was described earlier (Mir et al. 2018). Sheep polyclonal antibody detecting total S6K1 was purified at the University of Dundee (S417B). The anti-phospho-S6K1 Thr389 and the anti-phospho-MYPT1 Thr853 antibodies were purchased from Cell Signalling Technology (#9205 and #4563, respectively). For immunoblotting applications all commercial monoclonal antibodies were diluted in 5% (by vol) bovine serum albumin in TBS-T (20 mM Tris base, 150 mM Sodium Chloride (NaCl), 0.2% (by vol) Tween20), and sheep polyclonal antibodies were diluted in 5% (by vol) milk in TBS-T.

### Reagents

MLi-2 inhibitor was synthesized by Natalia Shpiro (University of Dundee) as described previously [27]. Calyculin A (ab141784) and Okadaic Acid (ab120375) were purchased from Abcam. Microcystin-LR was obtained from Enzo Life Sciences (ALX-350-012).

### Plasmids

The following cDNA constructs were used for transfections: HA-Empty (DU44059), HA-Rab10 (DU44250), HA-Rab8a (DU35414), HA-Rab12 (DU48963), HA-Rab3a (DU51539), HA-Rab35 (DU26478), HA-Rab43 (DU26392), HA-PPM1H (DU62789), PPM1H [H153D] (DU62928), PPM1H [D288A] (DU62985), HA-PPM1H D288E (DU62986), Flag-LRRK2 [R1441G] (DU13077).

These are available from MRC PPU Reagents and Services (https://mrcppureagents.dundee.ac.uk/). DNA constructs were amplified in Escherichia coli DH5α and purified using a Hi-Speed Plasmid Maxi Kit (Qiagen).

### siRNA Screens

The siRNA screen was performed using a human siRNA library (Dharmacon) designed to target 322 phosphatases. The list of siRNA targets and the sequences of all siRNA oligonucleotides used are provided in Supplementary Excel File 1. A549 cells were seeded in 6-well plates at 150,000 cells/well. After 24 h cells were transfected using 2 µl Lipofectamine RNAi Max and 20 pmol of siRNA per well. Cells were then cultured for a further 72 hours. In Screen 1 and 2, cells were directly lysed without further treatment, whereas in Screen 3, cells were treated for 5 min with 100 nM MLi-2 prior to lysing. Lysates were centrifuged at 20,800 g for 15 minutes at 4°C, quantified by Bradford assay (Thermo Scientific) and subjected to immunoblot analysis.

### Generation of PPM1H CRISPR/Cas9 knockout

CRISPR was performed using a paired nickase approach to minimize off-target effects [59]. Analysis of the PPM1H locus (ENSG00000111110) showed the expression of a single verified transcript (NM_020700.2, ENST00000228705.7) and exon 1 specific guide pairs with low combined off-targeting scores were subsequently identified using the Sanger Institute CRISPR webtool (http://www.sanger.ac.uk/htgt/wge/find_crisprs). Complementary oligos for the optimal guide pair A (G1 5’- gCAATTTCATGGGCGGCATCA and G2 5’- gCTCGAGTGAGCATATTACTC) were designed and annealed according to the Zhang method [60] with BbsI compatible overhangs facilitating cloning into the target vectors; the sense guide G1 was cloned into the puromycin selectable plasmid pBABED P U6 (DU48788, https://mrcppureagents.dundee.ac.uk/) and the antisense guide G2 cloned into the spCas9 D10A expressing vector pX335 (Addgene Plasmid #42335) yielding clones DU64249 and DU64253 respectively.

CRISPR was performed by co-transfecting A549 cells (80% confluency, 10cm dish) with 3.75 μg of each plasmid using 27 µl of Lipofectamine LTX according to the manufacturer’s instructions (Life Technologies). The transfected cells were then incubated for 24 h in DMEM supplemented with 10% (by vol) FBS, 2 mM L-glutamine, 100 units/ml penicillin and 100 μg/ml streptomycin. Following CRISPR, the medium was replaced with fresh medium supplemented with 3 μg/ml puromycin for a total of 48 hours to enrich for transfectants, adding fresh media each day. The cells were then grown for a further 48 hours in media without puromycin to provide sufficient cells for single-cell sorting.

Single cells were placed in individual wells of a 96-well plate containing DMEM supplemented with 10% (by vol) FBS, 2 mM L-glutamine, 100 units/ml penicillin, 100 μg/ml streptomycin and 100 μg/ml Normocin (InvivoGen). After reaching ∼80% confluency, individual clones were transferred into six-well plates and PPM1H expression determined via immunoblot. Wild type A549 cell were cultured side-by-side with the knock-out pools and subjected to single cell sorting. 2 independent wild-type clonal cell lines were obtained and used as controls for knock-out clones.

### Immunoprecipitation Assays

Flag-LRRK2 [R1441G] and HA-PPM1H, HA-PPM1H[D288A], or HA-empty were transiently overexpressed in HEK293 cells using Polyethylenimine transfection [61]. 24 h post-transfection, cells were lysed in lysis buffer as described above and HA-tagged proteins were immunoprecipitated using anti-HA beads overnight (40 µl resin for 2 mg protein). Immunoprecipitates were then washed twice with lysis buffer containing 0.5 M NaCl and a further three times with Dulbecco’s phosphate-buffered saline (200 mg/L Potassium Chloride (KCl), 200 mg/L Potassium Phosphate monobasic (KH2PO4), 8000 mg/L Sodium Chloride (NaCl), 2160 mg/L Sodium Phosphate dibasic (Na2HPO4-7H2O)). 40 µl lithium dodecyl sulfate loading buffer (106 mM Tris HCl, 141 mM Tris Base, 2% (by mass) LDS, 10% (by vol) Glycerol, 0.51 mM EDTA, 0.22 mM SERVA Blue G250, 0.175 mM Phenol Red, pH 8.5), diluted 2-fold and then added to the beads. The mixture was then incubated at 100°C for 10 min, and the eluent was collected by centrifugation through a 0.22-μm-pore-size Spin-X column and then supplemented with 2-Mercaptoethanol to a final concentration of 1% (by vol). Samples were incubated for 5 min at 70°C before being subjected to immunoblot analysis.

### In-gel digestion of PPM1H Immunoprecipitation

Wild type and mutant PPM1H was expressed in HEK293 cells, immunoprecipitated and immunoprecipitates washed as described above and elution performed by adding 40 µl of immunoprecipitate (50% slurry) to 40 µl of 4x LDS (106 mM Tris HCl, 141 mM Tris Base, 2% (by mass) LDS, 10% (by vol) Glycerol, 0.51 mM EDTA, 0.22 mM SERVA Blue G250, 0.175 mM Phenol Red, pH 8.5). 10% of the immunoprecipitate was kept back for immunoblot analysis with the remainder being reduced by adding dithiothreitol to 5 mM in a total volume of 50 µl and the resultant mixture incubated at 56oC for 20 min on a Thermomixer with 800 rpm agitation. The tubes were brought to room temperature and iodoacetamide added to a final concentration of 20 mM and the samples were incubated in dark at room temperature for 20 min on a Thermomixer with 800 rpm agitation. The samples were resolved on a 4-12% Bis-Tris gel (Invitrogen C USA, NP0321) which is subsequently stained with Colloidal Coomassie blue (Part no: LC6025., Invitrogen, CA, USA). After the distaining, the 20-30 kDa gel region containing Rab proteins were excised, and incubated with 200 µl of 40 mM ammonium bicarbonate in 40% acetonitrile for 20 min. The gel pieces were dehydrated by washing twice with 100 ul of 100% Acetonitrile by incubating at room temperature for 20 min. The Acetonitrile was removed, and the gel slices were rehydrated by adding 100 µl of 20 mM triethylammonium bicarbonate buffer pH7.5 containing 400 ng of trypsin. Samples were digested overnight by incubating on a Thermomixer at 37oC. The tryptic peptides were extracted by washing the gel pieces with 100 µl of 80% acetonitrile in 0.5% (by vol) acetic acid. This elution step was repeated twice, eluates combined, peptides vacuum dried and desalted using C18 stage tips as described previously [16].

### LC-MS/MS analysis

The desalted peptides were dissolved in 50 µl of solvent-A (0.1% formic acid in water) prior to loading on to Evotips (https://www.evosep.com/) as described in [62]. Briefly, the Evotips are activated by incubating in 100% isopropanol and then washed twice with 20 µl of Solvent-B (100% Acetonitrile in 0.1% formic acid by volume) and subsequently equilibrated by washing twice with 20 µl of solvent-A (0.1% formic acid in water). The acidified peptide digest was applied to the Evotips and washed twice with 20 µl of Solvent-A and an additional 100 µl solvent-A was added before placing them on auto sampler rack of EvoSep-ONE liquid chromatography system which is connected to Thermo Q Exactive HFX mass spectrometer (Thermo Fisher Scientific, Bremen, Germany) through Easy spray nano electrospray ionization source. As described in [62], EvoSep One LC system applies a partial elution by applying 35% Solvent-B (100% Acetonitrile in 0.1% formic acid) through the Evotip and subsequently the eluted peptides are pushed into a long storage capillary which is in line with a high pressure pump by a 6-port rotary valve. Following the partial elution, the high-pressure pump pushes the peptide sample into an 8cm analytical column (PN: EV-1074; 100um ID, 15cm long, C18 AQ, 3um beads., Dr Maisch, Ammerbuch, Germany) and electro sprayed directly into the mass spectrometer. The peptides were resolved by selecting 45 min run method and acquired in a data dependent mode on QE-HFX. Top 10 precursor peptides were selected in each survey scan between 350 to 1500 m/z. The MS1 scans were acquired at 120,000 resolution at m/z 200 and measured in Ultra high-field Orbitrap mass analyzer. The precursor ions were fragmented at 27 NCE using higher energy collisional dissociation and the product ion MS2 scans were acquired at 15,000 resolution at m/z 200 and measured in Ultra high-field Orbitrap mass analyzer. The maximum AGC target and Ion injection times were maintained at 3E6 and 50 ms for MS1 and 5E4 and 120 ms for MS2 scans. Singly charge and charge states above 5 are excluded and the dynamic exclusion was selected for 10 sec to avoid the repeated acquisition of previously acquired peptide precursors

### Data analysis and Label free quantification

The MaxQuant software suite [63] version 1.6.6.0 was used for database search with the following parameter. Standard search engine, Trypsin/P was selected as a specific enzyme by allowing 2 missed cleavages, Oxidation of (M), Acetyl (Protein-N-terminal), Deamidation N and Q were selected as variable modifications. Carbamidomethylation Cys was selected as fixed modification. LFQ option was enabled by selecting LFQ min ratio count as 2. First search tolerance of 20 ppm and main search tolerance of 4.5 ppm were selected. Global Parameters: Uniprot Human protein database was used for the database search and 1% protein, peptide and PSM FDR was enabled and match between runs was enabled within groups. For protein quantification, min ratio count was set at 2 for accurate LFQ quantification. The MaxQuant output protein group text files were processed using Perseus software suite [64], version 1.6.2.3 was used. The data was filtered for any proteins that are identified only by site, common contaminants and reverse hits and proteins identified with single unique peptides. Following, the LFQ intensities were log2 transformed and Student-T test and permutation-based FDR was applied to identify the differentially enriched and significant protein groups between HA-PPM1H-D288A and Wild type HA-PPM1H.

### Expression and purification of recombinant PPM1H

4 x 500 ml of Lysogeny broth containing 100 µg/ml ampicillin antibiotic was inoculated with a single colony of BL21-CodonPlus(DE3)-RIPL strain of E. coli transformed with either Plasmid DU62835 (expresses His-SUMO-PPM1H[wildtype]), Plasmid DU68104 (His-SUMO-PPM1H[H153D]), or Plasmid DU68087 (His-SUMO-PPM1H[D288A]). Bacteria were cultured at 37oC until OD600 is 0.4-0.6. Temperature was reduced to 15oC and protein expression was induced by addition of Isopropyl β-D-1-thiogalactopyranoside to 50 µM in addition to MnCl2 to 2 mM as PPM family of phosphatases require Mn or Mg for stability [43]. Cells were cultured for 16 h before harvesting by centrifugation at 4,200 x g for 20 mins at 4oC. The pellet is resuspended in 200 ml of ice cold E.coli lysis buffer [50 mM Tris/HCl pH7.5, 150 mM NaCl, 1% (by vol) Triton, 2 mM MnCl2, 0.5 mM TCEP (tris(2-carboxyethyl)phosphine)), 1 mM Pefabloc (4-(2-aminoethyl)-benzene-sulfonyl fluoride) and 1 mM benzamidine. Cells were lysed using a cell disruptor (passing sample through twice) and extracts clarified by centrifugation at 30,000 x g for 20 min at 4oC. Lysates were incubated in 2 ml of Cobalt-Agarose (Amintra Cobalt NTA Affinity Resin, Expedeon) that has been equilibrated in E. coli lysis buffer and incubate on roller at 4°C for 90 minutes. The resin was loaded onto a column and wash with 20 column volumes of High Salt Wash Buffer [50 mM Tris/ HCl pH7.5, 500 mM NaCl, 2 mM MnCl2, 0.03% (by vol) Brij 35, 20 mM Imidazole, 0.5 mM TCEP] until no unbound protein is present in the flow-through. The column was wash with 5 column volumes of Low Salt Wash buffer [50 mM Tris/ HCl pH7.5, 150 mM NaCl, 2 mM MnCl2, 10 mM Imidazole, 0.03% (by vol) Brij 35, 0.5 mM TCEP]. Protein was eluted with Elution Buffer [Low Salt Wash + 500 mM imidazole pH7.5] and 1 ml fractions collected. The fractions containing protein were pooled and concentrated to 1 ml final volume using an Amicon Ultra-15 30 kDa concentrator (Millipore, Z717185). This was subjected to gel filtration purification on a Superdex 75 Increase 10/300GL column equilibrated in Equilibration Buffer [50 mM Tris/HCl pH7.5, 150 mM NaCl, 2 mM MnCl2, 0.5 mM TCEP], collecting 0.2 ml fractions at a flow rate of 0.2 ml/min. Fractions containing wild type and mutant His-Sumo-PPM1H eluted at the expected molecular weight region for (68 kDa protein) were pooled and dialyzed into Equilibration Buffer containing 270 mM Sucrose and stored in aliquots at -80oC.

### Expression and purification of Rab8A

Rab8A residues 1-181 was produced in 6 versions: the Rab8A [Q67L] mutant in the presence of 1 µM GTP-gamma-S and the wildtype in the presence of 10 µM GDP or in the presence of 5 µM GTP-gamma-S. All three were made as unphosphorylated and phosphorylated proteins.

Rab8A 1-181 [Q67L] was produced in the presence of GTP-gamma-S as follows. The codon optimized coding sequence of Rab8A residues 1-181 [Q67L] subcloned into pET28b was kindly provided by Amir Khan (Trinity College Dublin) and expressed based on a protocol provided by the Khan lab. Briefly, the expression construct was transformed into BL21 DE3 and expressed in Lucia Broth medium (Merck, Darmstadt) supplemented with 50 µg/ml Kanamycin (Formedium, UK). Typically, 12 x 1L expressions were set up. When the optical density at 600 nm reached 0.6, the culture temperature was dropped to 18oC and after 60 min, Rab8A expression was induced by addition of Isopropyl β-D-1-thiogalactopyranoside (Formedium, UK) to 0.1 mM for incubation overnight. The cells were collected by sedimentation and resuspended in ice cold lysis buffer (50 mM Tris pH 7.5, 10% (by vol) glycerol, 250 mM NaCl, 15 mM Imidazole, 2 mM MgCl2, 0.4% (by vol) Triton X-100, 1 mM Pefabloc, 10 µg/ml Leupeptin) employing 20 ml Lysis buffer per 5 ml of cell sediment. The cell suspension was sonicated, and insoluble material was removed by centrifugation (35000 x g at 4oC for 30 min). The supernatant was incubated with 3 ml Ni-agarose (Expedion, UK) for 90 min, which was then washed 5 times with 10 vol of wash buffer (50 mM Tris pH 7.5, 5% (by vol) glycerol, 250 mM NaCl, 15 mM Imidazole, 14 mM 2-mercaptoethanol, 2 mM MgCl2, 0.03% (by vol) Brij35, 1 µM GTP-gamma-S). The protein was eluted with 4 x 1 resin volume of elution buffer (30 mM Tris pH 7.5, 3% (by vol) glycerol, 0.4 M imidazole, 8 mM 2-mercaptoethanol, 1.2 mM MgCl2, 0.02% (by vol) Brij35, 0.6 µM GTP-gamma-S) and dialyzed against 50 mM Tris pH 7.5, 10% (by vol) glycerol, 250 mM NaCl, 14 mM 2-mercaptoethanol, 2 mM MgCl2, 0.6 µM GTP-gamma-S in the presence of 60 Units of Thrombin (Sigma-Aldrich T1063-1KU). The sample was then passed through a Ni-agarose column (3 ml) equilibrated in 50 mM Tris pH 7.5, 10% (by vol) glycerol, 250 mM NaCl, 14 mM 2-mercaptoethanol, 2 mM MgCl2 and 0.6 µM GTP-gamma-S. The protein that was eluted (typically 25 mg) was subjected to chromatography on a Superdex-75 XK 26/60 column equilibrated in 50 mM Tris pH 7.5, 10% (by vol) glycerol, 250 mM NaCl, 14 mM 2-mercaptoethanol, 2 mM MgCl2 and 0.6 µM GTP-gamma-S, collecting 1.5 ml fractions at a flow rate of 2.5 ml/min. Fractions containing Rab8A eluted at the expected molecular weight region for (20 kDa protein) were either concentrated to above 1 mg/ml to be used as unphosphorylated Rab8A or - for the production of phospho-Rab8A - directly phosphorylated, avoiding both concentration and freeze thawing to mitigate aggregation. Up to 16 mg of Rab8A could be produced from a 12 Liter preparation.

Wild type Rab8A 1-181 (wildtype, truncated) was produced either in the presence of 10 µM GDP or in the presence of 5 µM GTP-gamma-S respectively present in all solutions throughout. Otherwise the expression and purification method was identical to the method used for the purification of Rab8A 1-181 [Q67L].

### Expression and Purification of Rab10

The method for expressing and purifying Rab10[1-181] (GDP-bound) was similar to the method for purifying GDP-bound Rab8A. Rab10 is much more aggregation prone than Rab8A and we found that the addition of 20 mM L-arginine and 0.5M NaCl in the lysis buffer, wash buffers and dialysis buffer was used to mitigate aggregation. Yield was at over 10 times lower than with Rab8A.

### Generation of Thr72 stoichiometrically phosphorylated Rab8A (pRab8A)

It has recently been shown that the MST3 kinase can specifically and efficiently phosphorylate Rab8A at Thr72 in vitro (Vieweg, S et al manuscript in preparation). As MST3 is much easier to express than LRRK2, we decided to phosphorylate Rab8A at Thr72 using recombinant MST3. 6His-MST3 residues 1-431 (DU62878) was obtained from MRC-PPU Reagents and Services (https://mrcppureagents.dundee.ac.uk/reagents-proteins/overview). 5 mg of His-MST3 were incubated with 12 mg Rab8A in a total volume of 20 ml in Equilibration buffer, supplemented with 10 mM MgCl2 and 1 mM ATP at 27oC overnight. 6His-MST3 was removed from the reaction by passing the reaction through an 3 ml Ni-agarose column equilibrated in 50 mM Tris pH 7.5, 5% (by vol) glycerol, 250 mM NaCl, 14 mM 2-mercaptoethanol, 15 mM Imidazole, 2 mM MgCl2, 0.03% (by vol) Brij35, 1 µM GTP-gamma-S). The phosphorylated Rab8A sample was diluted 7-fold into Source-S Buffer [20 mM MES pH 5.3, 10 mM NaCl, 10% (by vol) glycerol, 0.03% (by vol) Brij 35, 14 mM 2-mercaptoethanol, 2 mM MgCl2, 1 µM GTP gamma-S. The phospho-Rab8A was separated from the unphosphorylated Rab8A by chromatography over a Source 15 S HR10/10 column. The column was equilibrated and run in Source-S Buffer at a flowrate of 2 ml/min and a gradient from 10 mM NaCl to 0.5M NaCl deployed and 1 ml fractions were collected. Phosphorylated Rab8A eluted at around 10 mS/cm (∼80-110 mM NaCl), whereas a much smaller amount of non-phosphorylated Rab8A eluted after 15 mS/cm (Figure 7-Figure Supplement 1). Analysis of the phosphorylated and unphosphorylated Rab8A on 12% Phos-tag gels (Figure 7-Figure Supplement 1B) and mass spectrometry (Figure 7-Figure Supplement 2) confirmed that the first eluting peak was only significantly phosphorylated at Thr72 and the stoichiometry of phosphorylation was judged to be ∼97%. The fractions containing the phosphorylated Rab8A were concentrated, pooled, aliquoted, snap frozen in liquid nitrogen and stored at -80oC. GDP bound Rab8A was produced by supplementing all solutions during the preparation with 10µM GDP / 10mM MgCl2. Likewise, GTPgammaS Rab8A was produced in the presence of 2-5µM GTPgammaS / 10mM MgCl2 throughout the preparation.

### Generation of phosphorylated Rab10 protein

2.2mg of monomeric, pure Rab10 residues 1-181 was incubated with 0.5mg His-MST3 in the presence of 10mM MgCl2 and 1mM ATP at 27°C for 16h. The kinase was removed by depletion over a 0.2ml Ni-agarose bed. The flow-through was diluted 15-fold into 20 mM MES pH 5.3, 10 mM NaCl, 10% (by vol) glycerol, 0.03% (by vol) Brij 35, 14 mM 2-mercaptoethanol, 2 mM MgCl2, 10µM GDP and purified over a 10ml Source 15 S column as described for Rab8A [Q67L]. The column was developed with a NaCl gradient, here in the presence of 10µM GDP. Recovery was poor, but 0.1mg of phospho Rab10 could be obtained. Phospho Rab10 eluted at 14mS/cm and unphosphorylated Rab10 at 15 and 17 mS/cm respectively (Figure 7-Figure Supplement 4).

### PPM1H phosphatase assays employing pRab8A as substrate

In vitro dephosphorylation assay was performed in a total volume of 20 µl in Hepes buffer (pH 7.0) containing 10 mM MgCl2 using 2.5 µg pT72 phosphorylated Rab8a[Q67L] and varying levels of recombinant PPM1H. 3 µl of pT72 phosphorylated Rab8a[Q67L] at 0.83 mg/ml in a buffer containing 20 mM MES pH 5.3, 0.1 M NaCl, 10% (by vol) glycerol, 0.03% (by vol) Brij 35, 14 mM 2-mercaptoethanol, 2 mM MgCl2, 1 µM GTP-γ-S added to the assay. The assay was initiated by addition of 2.2 µl of serial dilutions of PPM1H (0.09 mg/mL, 0.018 mg/mL, 0.0036 mg/mL, 0.00072 mg/mL) diluted into HEPES Buffer from a stock of 0.45 mg/ml PPM1H in 50 mM Tris/HCl pH7.5, 150 mM NaCl, 2 mM MnCl2, 0.5 mM TCEP Buffer. The assay was carried out for 30 min and terminated by addition of 17.5 µl 4 x LDS (106 mM Tris HCl, 141 mM Tris Base, 2% (by mass) LDS, 10% (by vol) Glycerol, 0.51 mM EDTA, 0.22 mM SERVA Blue G250, 0.175 mM Phenol Red, pH 8.5) with 5 % (by vol) 2-Mercaptoethanol. Samples were then subjected to Phos-tag gel electrophoresis to determine stoichiometry of phosphorylated Rab8a[Q67L] as described previously [18]. Gel was stained using Instant Blue Coomassie (Expedeon).

### Overexpression of GFP-Rab10 and PPM1H in MEF cells

MEF cells stably expressing GFP-Rab10 were transiently transfected with plasmids carrying wild type HA-PPM1H, HA-PPM1H[H153D], or HA-PPM1H[D288A]. After 24 h the cells were fixed with 4% (by vol) PFA for 10 min, permeabilized with 0.1% saponin for 15 min, and blocked with 1% (by mass) BSA for 1 h. Cells were subsequently stained with mouse anti-HA antibody 1:1000 (Sigma-Aldrich H3663) and rabbit pRab10 1:1000 (Abcam ab230261). Highly cross absorbed H+L secondary antibodies (Life Technologies) conjugated to Alexa 568 or Alexa 647 were used at 1:5000. Primary and secondary antibody incubations were for 1 h at room temperature. All images were obtained using a spinning disk confocal microscope (Yokogawa) with an electron multiplying charge coupled device (EMCCD) camera (Andor, UK) and a 20x1.4NA or 40x1.4NA objective. Images were analyzed using CellProfiler and presented as maximum intensity projections. Results were quantified by determining the ratio numbers of pRab10 signal to GFP-Rab10 in cells expressing wild type HA-PPM1H, HA-PPM1H[H153D], or HA-PPM1H[D288A], and normalizing these numbers to the pRab10/GFP-Rab10 ratio in non-expressing cells. For each condition at least 20 cells were analyzed. Significance was determined by one-way analysis of variance with Dunnett’s post-test at 95% confidence interval. ***, P < 0.001.

### Immunofluorescence microscopy

RPE cells were transiently transfected with HA-PPM1H plasmid. After 24h, cells were treated for 2hr with 20 µM nocodazole in DMSO or DMSO alone, fixed with 4% (by vol) paraformaldehyde for 10 min, permeabilized with 0.1% by vol Triton X-100 for 5 min, and blocked with 1% (by mass) BSA for 1 h. Cells were subsequently stained with mouse or rabbit anti-HA antibody (Sigma-Aldrich H3663 or H6908, 1:1000) and the following Golgi markers: rabbit anti-GCC185 1:1000 (serum), rabbit anti-ß-Cop 1:1000 (serum), mouse anti-p1151:1000 (ascites), mouse anti-cation independent mannose 6-phosphate receptor 1:1 (2G11 culture sup), and rabbit anti-ACBD3 1:1000 (Sigma-Aldrich HPA015594). Highly cross absorbed H+L secondary antibodies (Life Technologies) conjugated to Alexa 488 or Alexa 568 were used at 1:5000. Primary and secondary antibody incubations were for 1 h at room temperature. Nuclei were stained with 0.1 µg/ml DAPI (Sigma). All images were obtained using a spinning disk confocal microscope (Yokogawa) with an electron multiplying charge coupled device (EMCCD) camera (Andor, UK) and a 100x1.4NA oil immersion objective. Images were analyzed using CellProfiler software [65] and presented as maximum intensity projections.

MEF cells stably expressing GFP-Rab10 were transiently transfected with plasmids carrying HA-PPM1H, HA-PPM1H H153D, or HA-PPM1H D288A. After 24 h the cells were fixed with 4% PFA for 10 min, permeabilized with 0.1% (by mass) saponin for 15 min, and blocked with 1% (by mass) BSA for 1 h. Cells were subsequently stained with mouse anti-HA antibody 1:1000 (Sigma-Aldrich H3663) and rabbit pRab10 1:1000 (Abcam ab230261). Highly cross absorbed H+L secondary antibodies (Life Technologies) conjugated to Alexa 568 or Alexa 647 were used at 1:5000. Significance was determined by one-way analysis of variance with Dunnett’s post-test at 95% confidence interval. ***, P < 0.001.

### Ciliation

MEF cells were infected in two rounds (day 1 and day 3) with lentiviruses encoding the shRNA sequences and on day 5, infected cells were selected using puromycin for 72 h as described [20]; pools of stably infected cells were then assayed for cilia formation. Ciliation was monitored after 24h serum starvation using anti-Arl13B antibody (NeuroMab, Davis, California) to stain cilia for immunofluorescence microscopy [20].

## Supporting information

Supplementary excel File 1 cited in text

Supplementary excel File 2 cited in text

## Acknowledgments

We thank Amir Kahn for providing us with the codon optimized Rab8a[1-181, 67QL], Ilaria Volpi for helping assembly of the phosphatase siRNA library, Miratul Muqit for providing the rabbit polyclonal Rab8A phospho-S111 antibody, Nicole K. Polinski, Marco Baptista and Shalini Padmanabhan (Michael J Fox Foundation for Parkinson’s research) for helpful discussions, the excellent technical support of the MRC-Protein Phosphorylation and Ubiquitylation Unit (PPU) DNA Sequencing Service (coordinated by Gary Hunter), the MRC-PPU tissue culture team (coordinated by Edwin Allen), MRC PPU Reagents and Services antibody and protein purification teams (coordinated by Hilary McLauchlan and James Hastie). This work was supported by the Michael J. Fox Foundation for Parkinson’s research [grant number 17298 (to S.R.P. and D.R.A.)] and [grant number 6986 (to S.R.P. and D.R.A.)] and MJFF Langston Award (to D.R.A which was used to purchase siRNA library); the Medical Research Council [grant number MC_UU_12016/2 (to D.R.A.)]; the pharmaceutical companies supporting the Division of Signal Transduction Therapy Unit (Boehringer-Ingelheim, GlaxoSmithKline, Merck KGaA -to D.R.A.) and the U.S. National Institutes of Health DK37332 (to S.R.P.).

## Author contributions

KB and PL designed and executed experiments in Figures 2, 3, 5, 6, 7, 8A and B.; WY designed and executed figure 10; PSW and WY designed and executed figures 4 and 9. FT designed and executed experiments in Figure 1; RN performed mass spectrometry for Figure 8C and expression analysis of PPM1H and PPM1M Fig 11-Figure Supplements 1 & 2. TM, generated expression constructs for CRISPR/CAS9 gene editing studies, MD expressed and purified PPM1H, PPM1M and PPM1J phosphatases, and MST3 kinase. AK expressed, purified and phosphorylated Rab8A for experiments shown in Figure 7, MW undertook most of the cloning; Suzanne R. Pfeffer and Dario R. Alessi supervised the project and wrote the manuscript. All authors were involved in discussing and interpreting the data.

---

**Figure 2—Figure Supplement 1.**
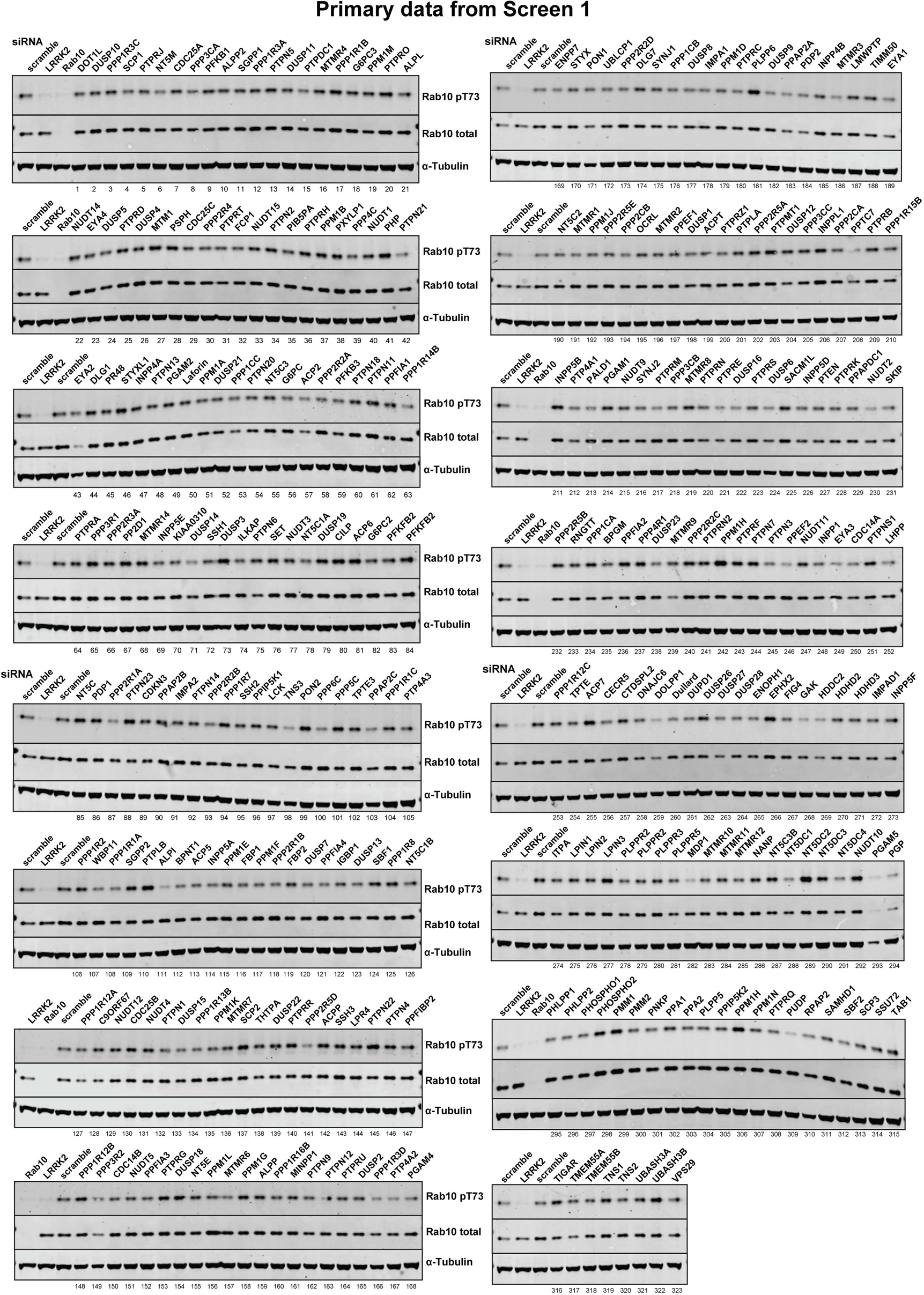
Primary data from siRNA Screen 1. Human A549 cells were transfected with the indicated siRNA pools (Dharmacon) for 72h. Cells were lysed and immunoblotted for the indicated antibodies and immunoblots developed using the LI-COR Odyssey CLx Western Blot imaging system. The numbering system corresponds to that employed in Figure 2 Supplement 1. The calculated intensities of the pRab10, Total Rab 10 and pRab10/Total Rab 10 ratio are provided in Supplementary Excel File 1

**Figure 2—Figure Supplement 2.**
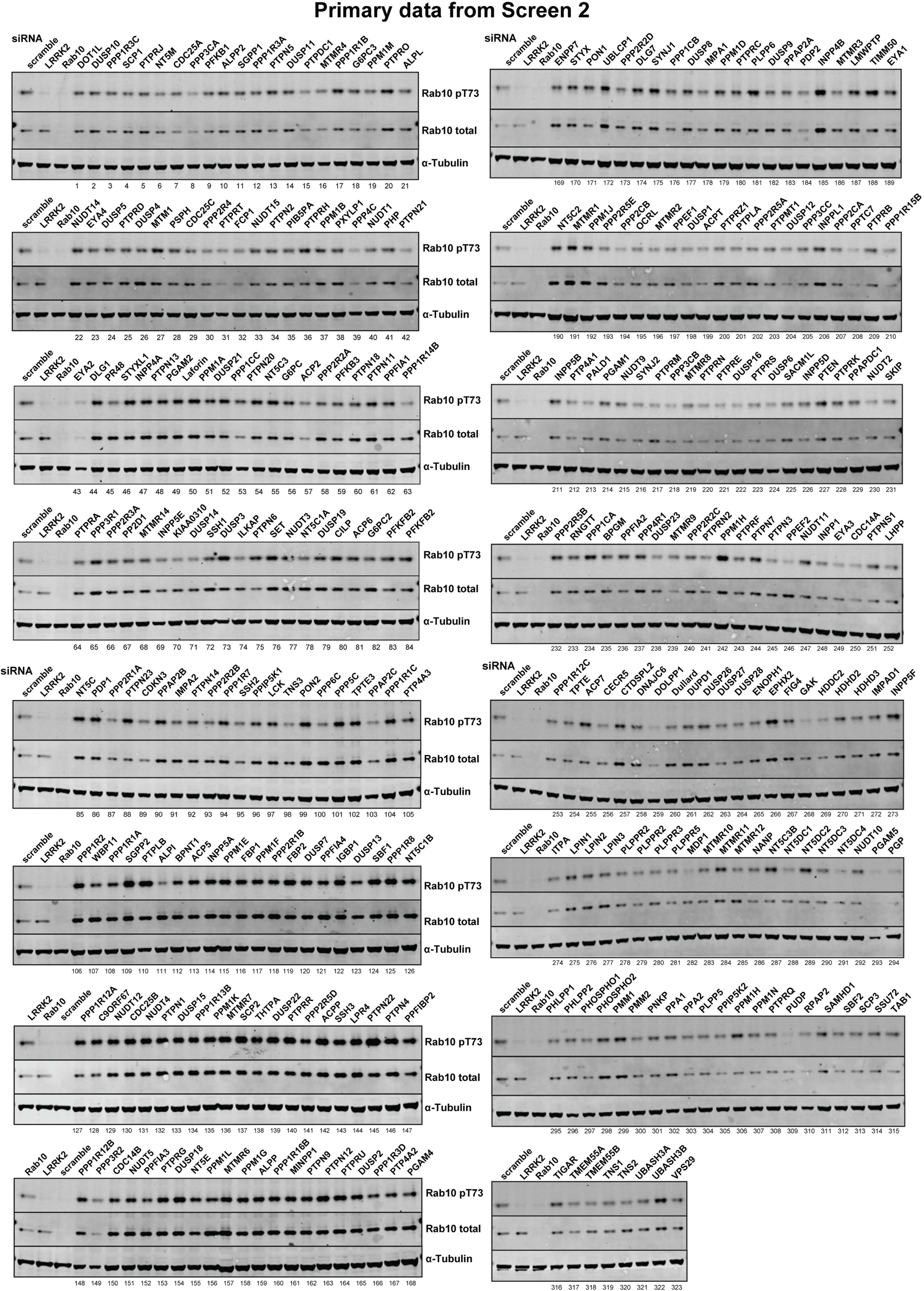
Primary data from siRNA Screen 2. Human A549 cells were transfected with the indicated siRNA pools (Dharmacon) for 72h. Cells were lysed and immunoblotted for the indicated antibodies and immunoblots developed using the LI-COR Odyssey CLx Western Blot imaging system. The numbering system corresponds to that employed in Figure 2 Supplement 1. The calculated intensities of the pRab10, Total Rab 10 and pRab10/Total Rab 10 ratio are provided in Supplementary Excel File 1

**Figure 2—Figure Supplement 3.**
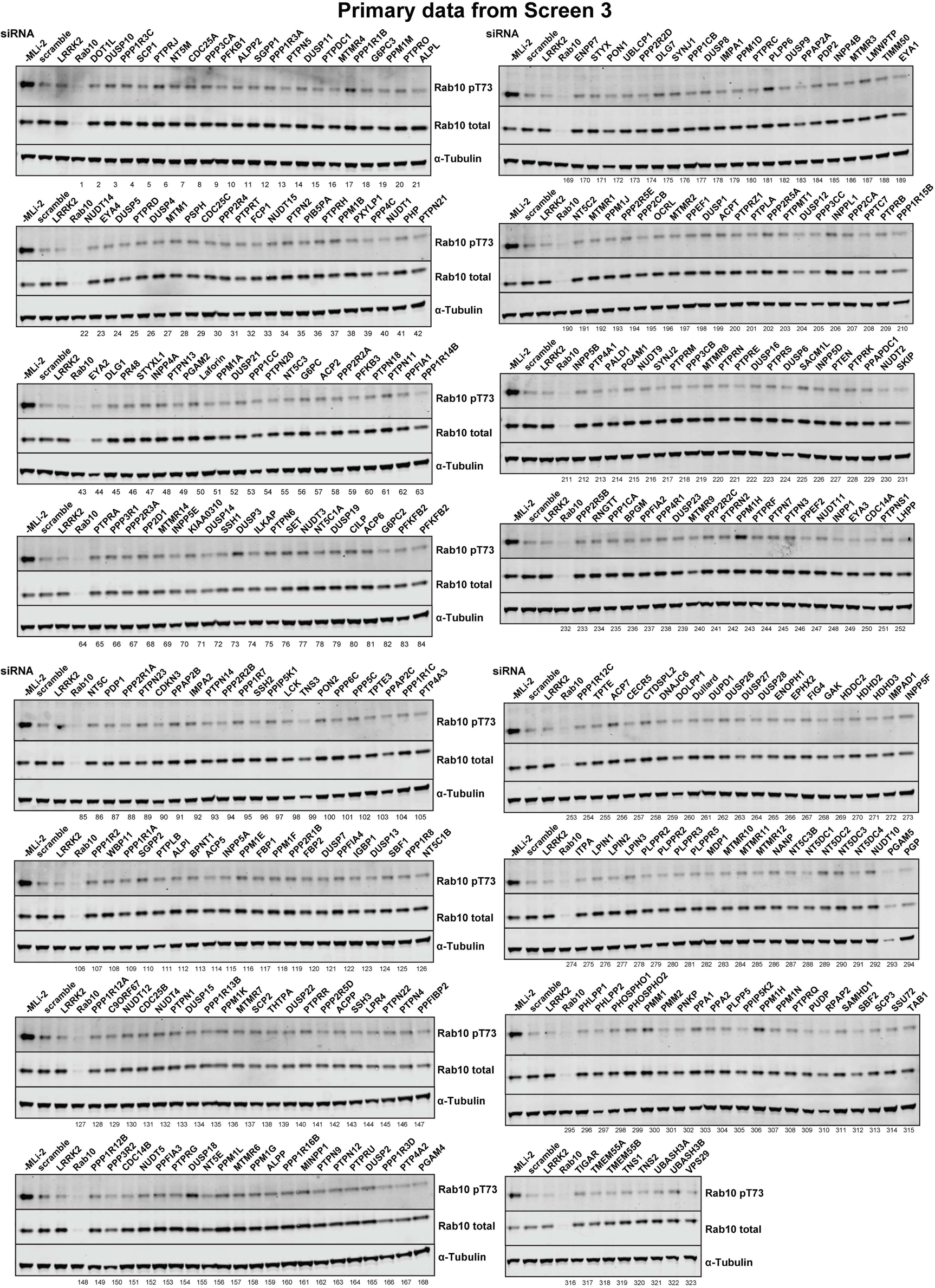
Primary data from siRNA Screen 3. Human A549 cells were transfected with the indicated siRNA pools (Dharmacon) for 72h. Cells were treated for 5 min with 100 nM MLi-2, lysed and immunoblotted for the indicated antibodies and immunoblots developed using the LI-COR Odyssey CLx Western Blot imaging system. The numbering system corresponds to that employed in Figure 2 Supplement 1. The calculated intensities of the pRab10, Total Rab 10 and pRab10/Total Rab 10 ratio are provided in Supplementary Excel File 1

**Figure 3-Figure Supplement 1.**
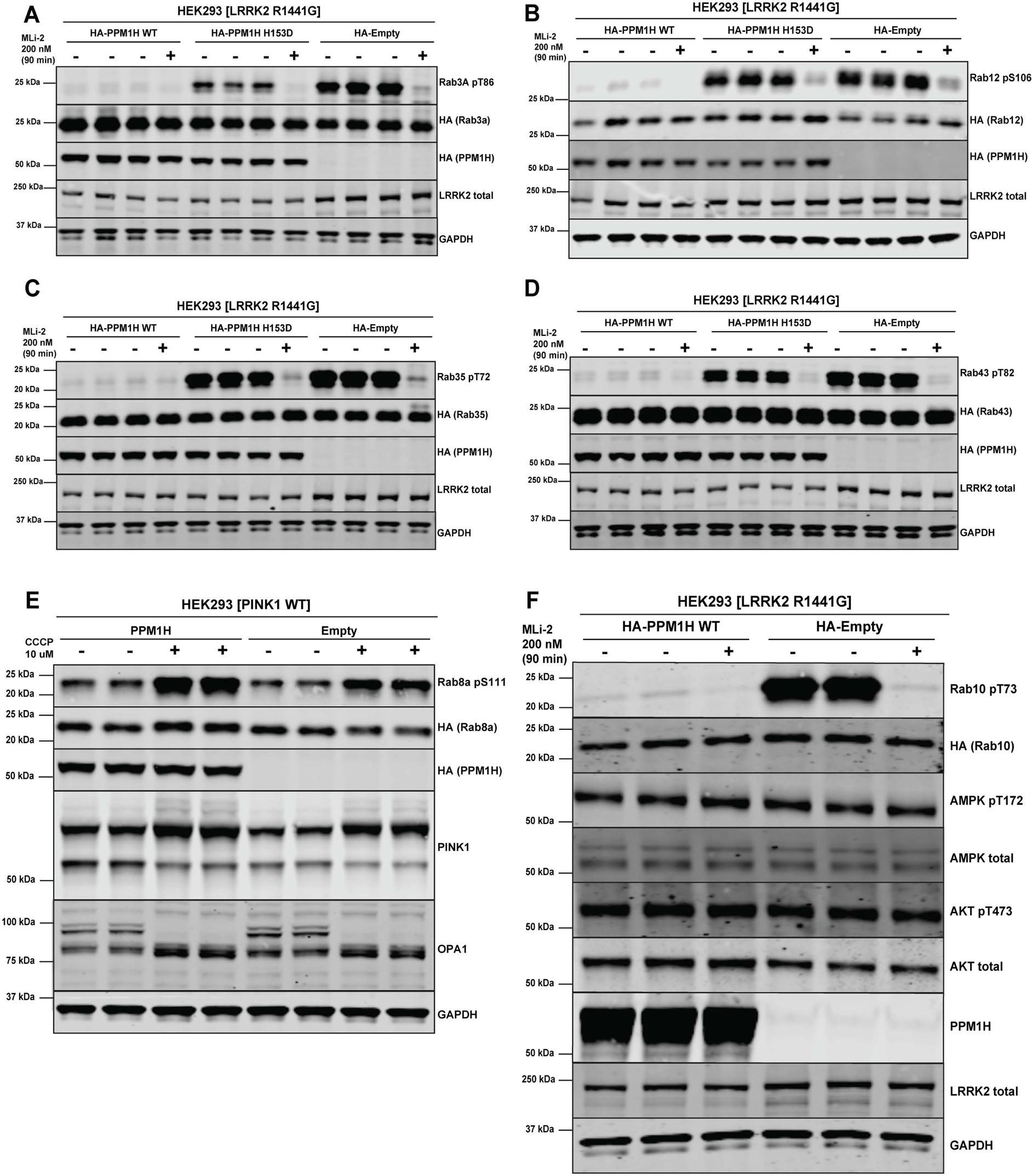
PPM1H dephosphorylates multiple LRRK2 phosphorylated Rab proteins. (A-D) HEK293 cells were transiently transfected with constructs expressing the indicated components. 24 h post-transfection, cells were treated with ± 200 nM MLi-2 for 90 min and then lysed. 10 µg whole cell lysate was subjected to LI-COR immunoblot analysis with the indicated antibodies at 1 µg/mL concentration. Each lane represents cell extract obtained from a different dish of cells. (E) As in (A) except that 24 h post-transfection, cells were treated ± 10 µM CCCP (Carbonyl cyanide m-chlorophenyl hydrazine) for 3 h to induce activation of the PINK1 kinase and trigger Rab8A phosphorylation at Ser111 [34]. (F) As in (A) except cells were immunoblotted with the indicated antibodies that recognize key phosphorylation sites of AMPK and Akt signalling pathway.

**Figure 5-Figure Supplement 1.**
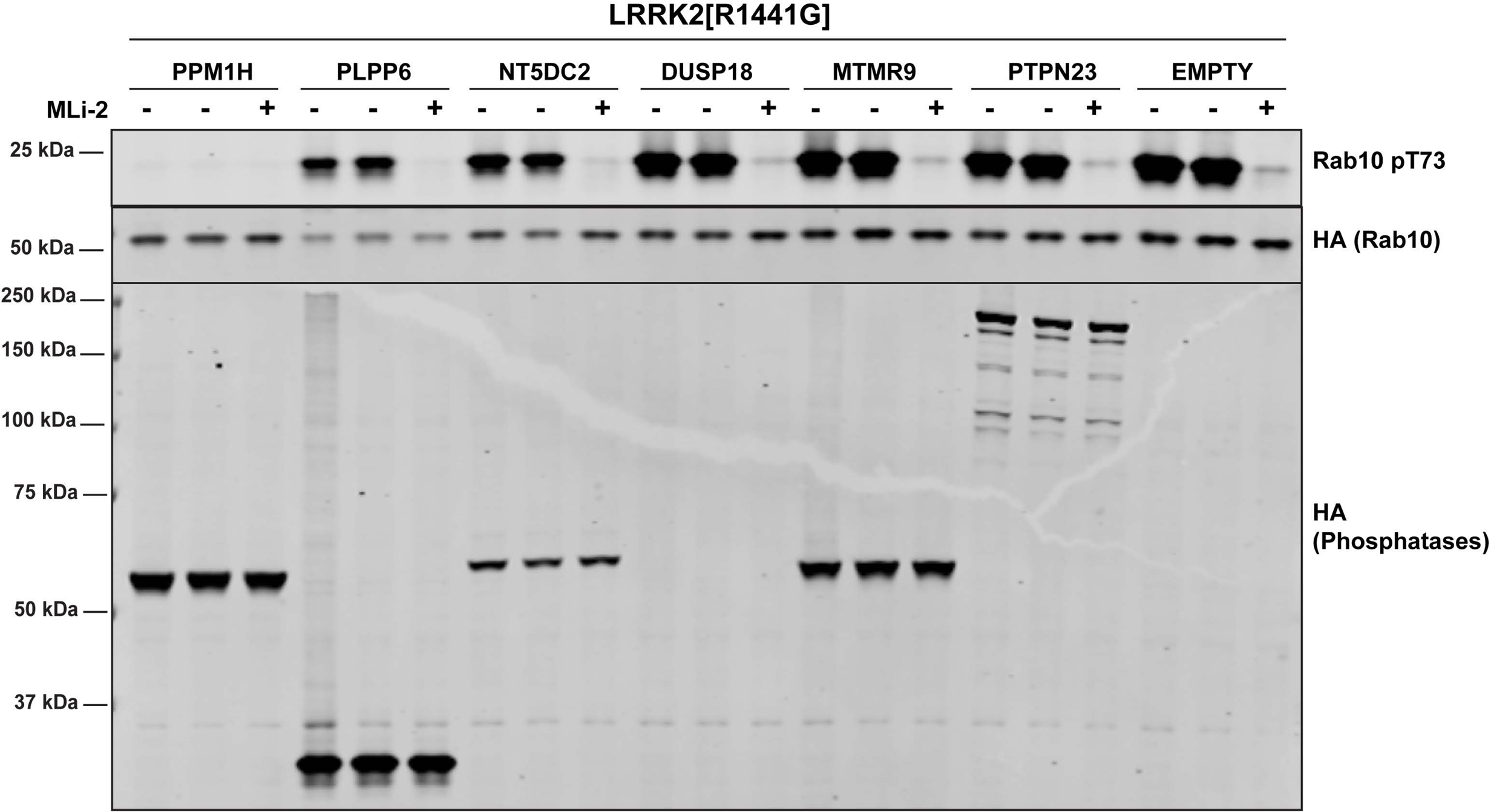
HEK293 cells were transiently transfected with constructs expressing the indicated constructs. 24 h post-transfection, cells were treated with ± 200 nM MLi-2 for 90 min and then lysed. 10 µg of whole cell lysate were subjected to immunoblot analysis with the indicated antibodies at 1 µg/mL concentration and membranes analyzed using OdysseyClx Western Blot imaging system. Each lane represents cell extract obtained from a different dish of cells.

**Figure 7-Figure Supplement 1.**
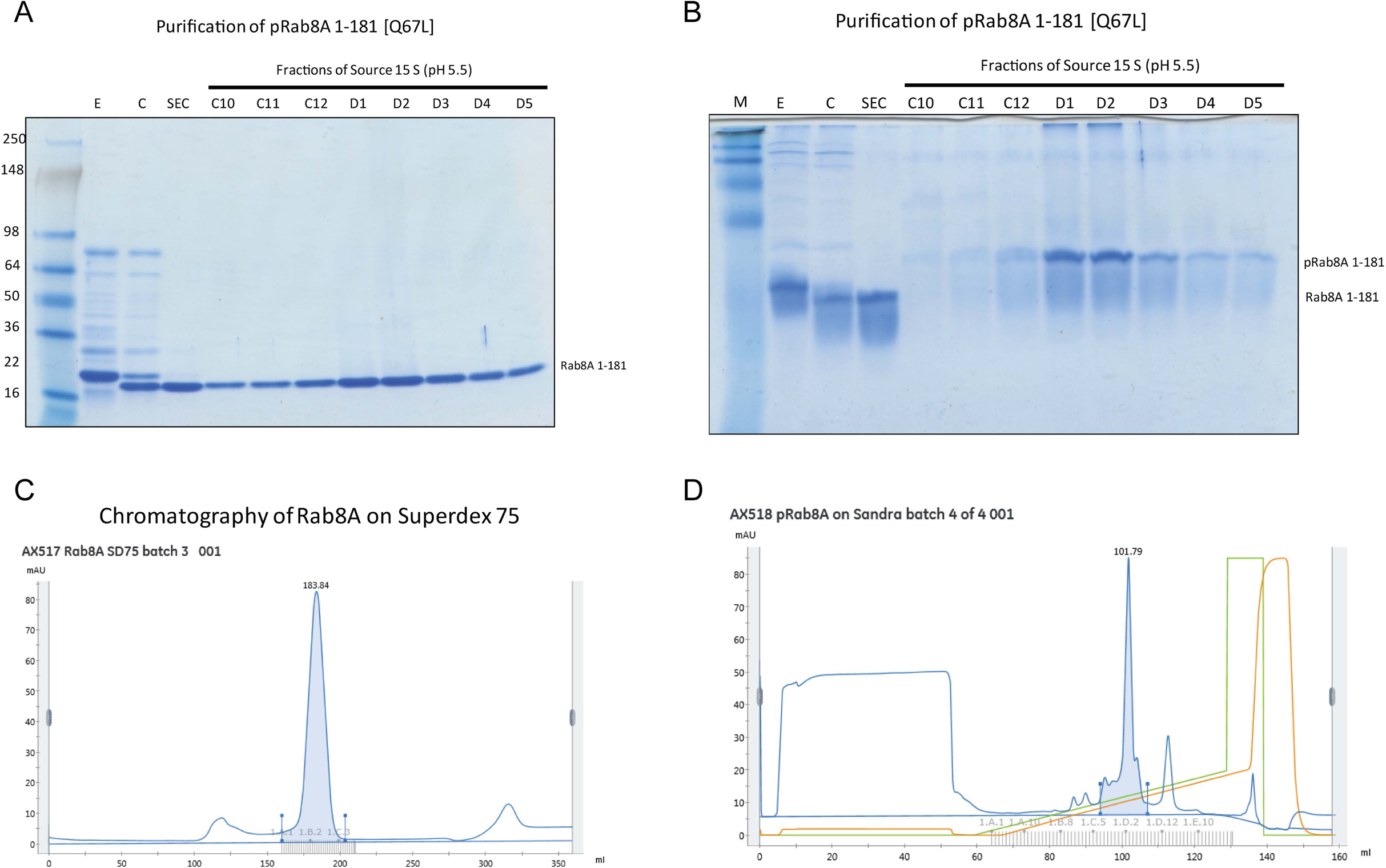
Purification and characterization of Thr72 phosphorylated Rab8A[1-181, Q67L]. (A) Coomassie stained 4-20% gradient SDS polyacrylamide gel electrophoresis analysis of the purification of pRab8A 1-181 [Q67L]. Lane M = See-Blue Plus Marker (Invitrogen); Lane E = His-Rab8A eluted from Ni-agarose. Lane C = Rab8A cleaved with Thrombin, dialyzed and depleted on Ni-agarose; Lane SEC = monomeric Rab8A after size exclusion chromatography; C10 – D5 = fractions of a 10 ml Source S column containing phospho-Rab8A 1-181 [[Q67L]. (B) Coomassie stained 12% Phos-tag SDS polyacrylamide gel electrophoresis analysis of the purification of pRab8A 1-181 [Q67L]. Lane M = See-Blue Plus Marker (Invitrogen); Lane E = His-Rab8A eluted from Ni-agarose. Lane C = Rab8A cleaved with Thrombin, dialyzed and depleted on Ni-agarose; Lane SEC = monomeric Rab8A after size exclusion chromatography; C10 – D5 = fractions of a 10 ml Source S column containing phospho-Rab8A 1-181 [[Q67L]. (C) Chromatogram (UV280 nm) of Rab8A 1-181, separated on a Superdex 75 XK 26/60 column. Equipment: Äkta Pure. Evaluation Software: Unicorn 7.1. Monomeric Rab8A eluted at 183.84 ± 10 ml. (D) Chromatogram (UV280nm) of Rab8A 1-181, separated on a Source 15 S HR10/10 column. Equipment: Äkta Pure. Evaluation Software: Unicorn 7.1. Phosphorylated Rab8A eluted at 101.79 ± 10 ml.

**Figure 7-Figure Supplement 2.**
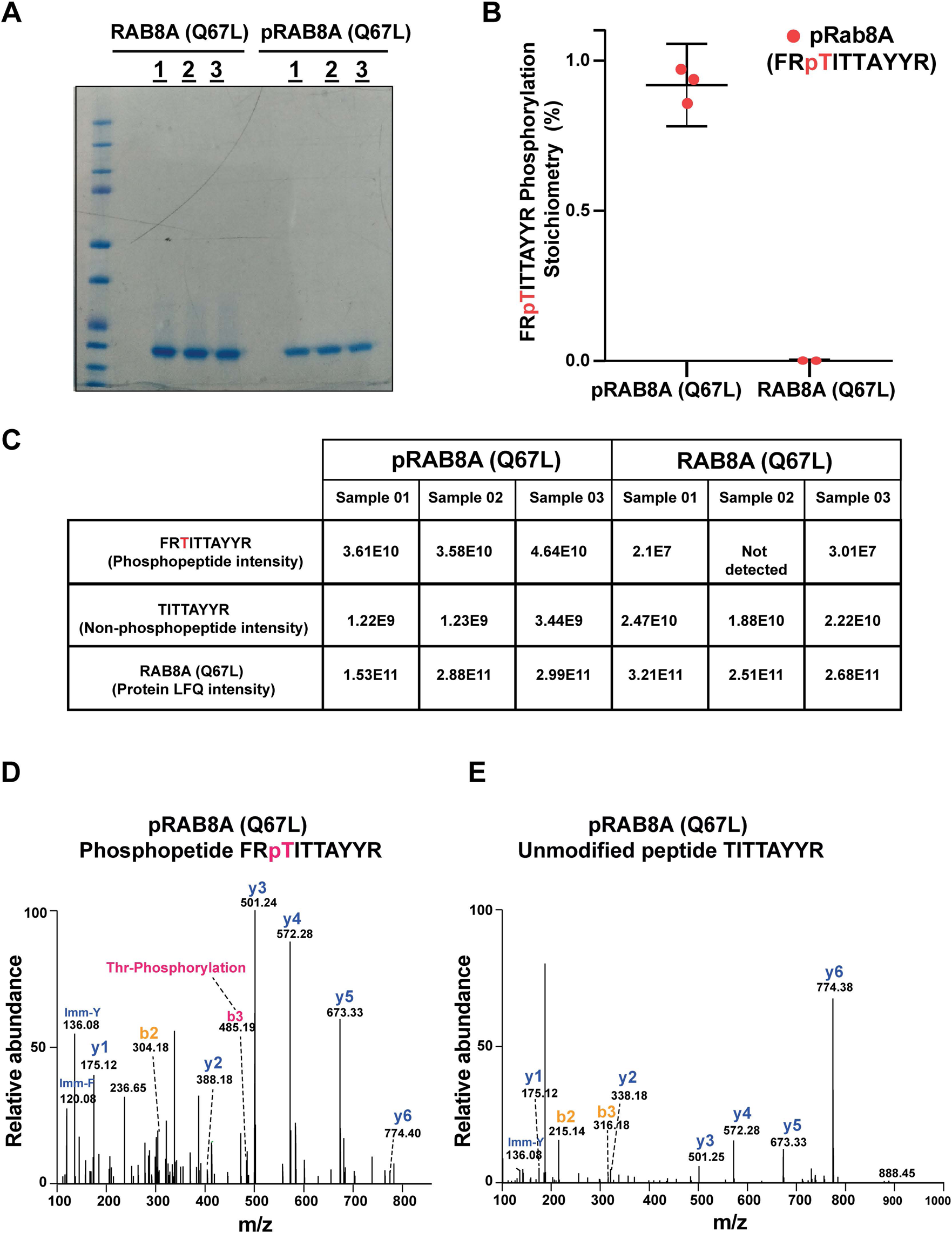
Mass spectrometry characterization of Thr72 phosphorylated Rab8A[1-181, Q67L]. (A) 2 µg of Rab8A 1-181 [Q67L] and pRab8A 1-181 [Q67L] in technical triplicates was loaded on a 4-12% Bis-Tris gradient gel. Following colloidal Coomassie staining, the lane encompassing 21 kDa region was excised and subjected to In-gel digestion using trypsin and analyzed on QE – HFX mass spectrometer. (B) and (C). The mass spectrometry raw data from Rab8A [1-181, Q67L] and pRab8A 1-181 [Q67L] samples were searched using MaxQuant pipeline. The Phosphopeptide FRpTITTAYR, unmodified peptide TITAYR intensities and pRab8A 1-181 [Q67L] and Rab8A 1-181 [Q67L] protein LFQ intensities as shown in table (C) were used to determine the FRpTITTAYR, phosphorylation stoichiometry To enable accurate phosphosite stoichiometry to be calculated both the modified and unmodified Thr72 peptides were excluded in determining the protein LFQ intensities. The phosphorylation stoichiometry in pRab8A 1-181 [Q67L] was found to be ∼94% and in Rab8A 1-181 [Q67L] was found to be <0.07% which is shown as separated scatter plot (B), the error bars represent the mean with 95% CI. (D) Representative annotated MS/MS spectrum of a Thr72, FRpTITTAYR (m/z 457.88587; Z=3) identified in pRAB8A 1-181[Q67L]. The m/z 485.19, z=1 of a b3 ion indicated with dotted lines confirms the phosphosite localization of the Phosphopeptide FRpTITTAYR. (E) Representative annotated MS/MS spectrum of an unmodified peptide, TITAYR (m/z 494.75793; z=2) identified in RAB8A 1-181 [Q67L].

**Figure 7-Figure Supplement 3.**
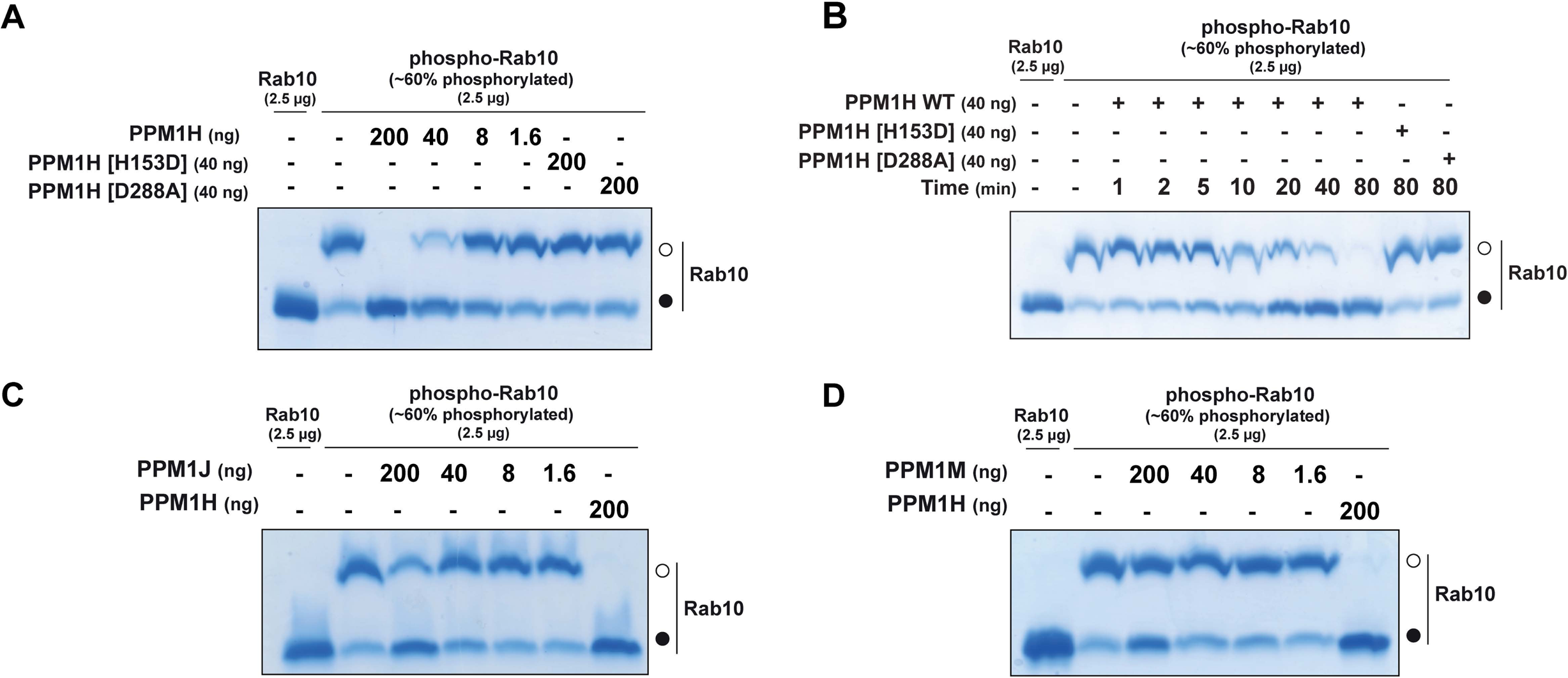
PPM1H dephosphorylates Rab10 in vitro. (A) The indicated amounts of recombinant wild type and mutant PPM1H (with a His-Sumo N-terminal tag, expressed in E.coli) were incubated in vitro with 2.5 µg pT73 ∼60% phosphorylated Rab10[1-181, GDP bound] for 30 min in the presence of 10 mM MgCl2 in HEPES pH 7.0 buffer. Reactions were terminated by addition of SDS Sample Buffer and analyzed by phos-tag gel electrophoresis that separates phosphorylated and dephosphorylated Rab10. Gel was stained with Instant Blue Coomassie. Bands corresponding to phosphorylated and non-phosphorylated Rab10 are marked with open (○) and closed (●) circles respectively. (B) As in (A) except that a time-course assay was performed using 2.5 µg pT73 phosphorylated Rab10[1-181, GDP bound] and 40 ng wild type or mutant PPM1H for the indicated times. (C) As in (A) except that PPM1J was assessed. (D) As in (A) except and PPM1M was assessed.

**Figure 7-Figure Supplement 4.**
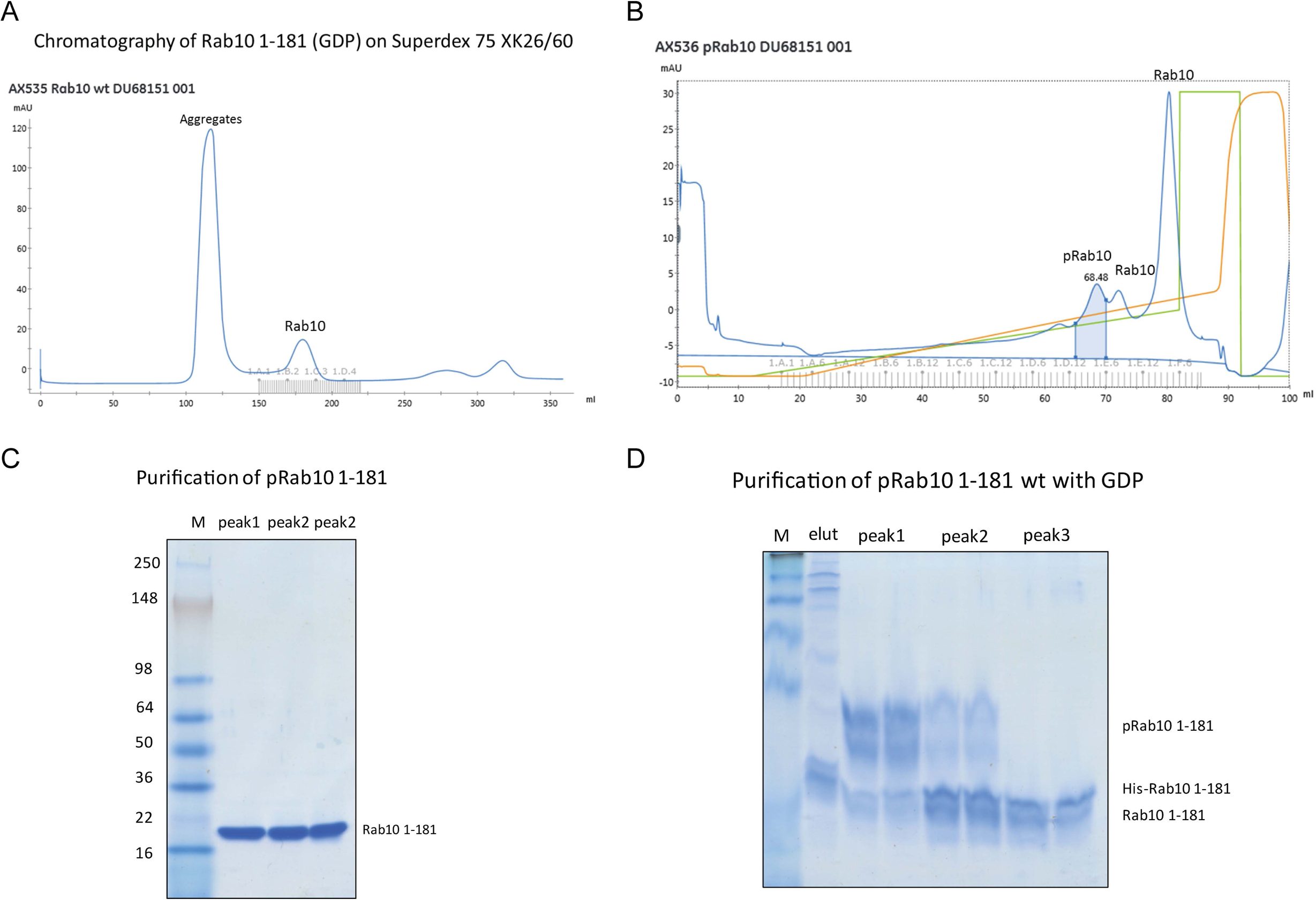
Purification and characterization of Thr73 phosphorylated Rab10[1-181]. (A) Chromatogram (UV280nm) of Rab10 1-181, separated on a Superdex 75 XK 26/60 column to obtain monomeric protein. Equipment: Äkta Pure. Evaluation Software: Unicorn 7.1. Monomeric Rab10 eluted at 180.03 ± 10ml. (B) Chromatogram (UV280nm) of phosphorylated Rab10 1-181 (wild type, wt with GDP), separated on a Source 15 S HR10/10 column. Equipment: Äkta Pure. Evaluation Software: Unicorn 7.1. Phospho-Rab10 eluted at 68.48 ± 2ml. (C) Coomassie stained 4-20% gradient SDS polyacrylamide gel electrophoresis analysis of the purification of pRab10 1-181 (wt in GDP). Lane M = See-Blue Plus Marker (Invitrogen); peak1 = first peak from a 10ml Source S column containing phospho Rab10 1-181. Peak2 = second peak from a 10ml Source S column containing Rab10. Peak3 = third peak, containing Rab10. (D) Coomassie stained 12% Phos-tag SDS polyacrylamide gel electrophoresis analysis of the purification of pRab10 1-181 (with GDP). Lane M = See-Blue Plus Marker (Invitrogen); Lane elut= His-Rab10 eluted from Ni-agarose ; peak1 = first peak from a 10ml Source S column containing phospho Rab10 1-181. Peak2 = second peak from a 10ml Source S column containing mostly Rab10. peak3 = third and major peak of the Source S column (unphosphorylated Rab10).

**Figure 7-Figure Supplement 5.**
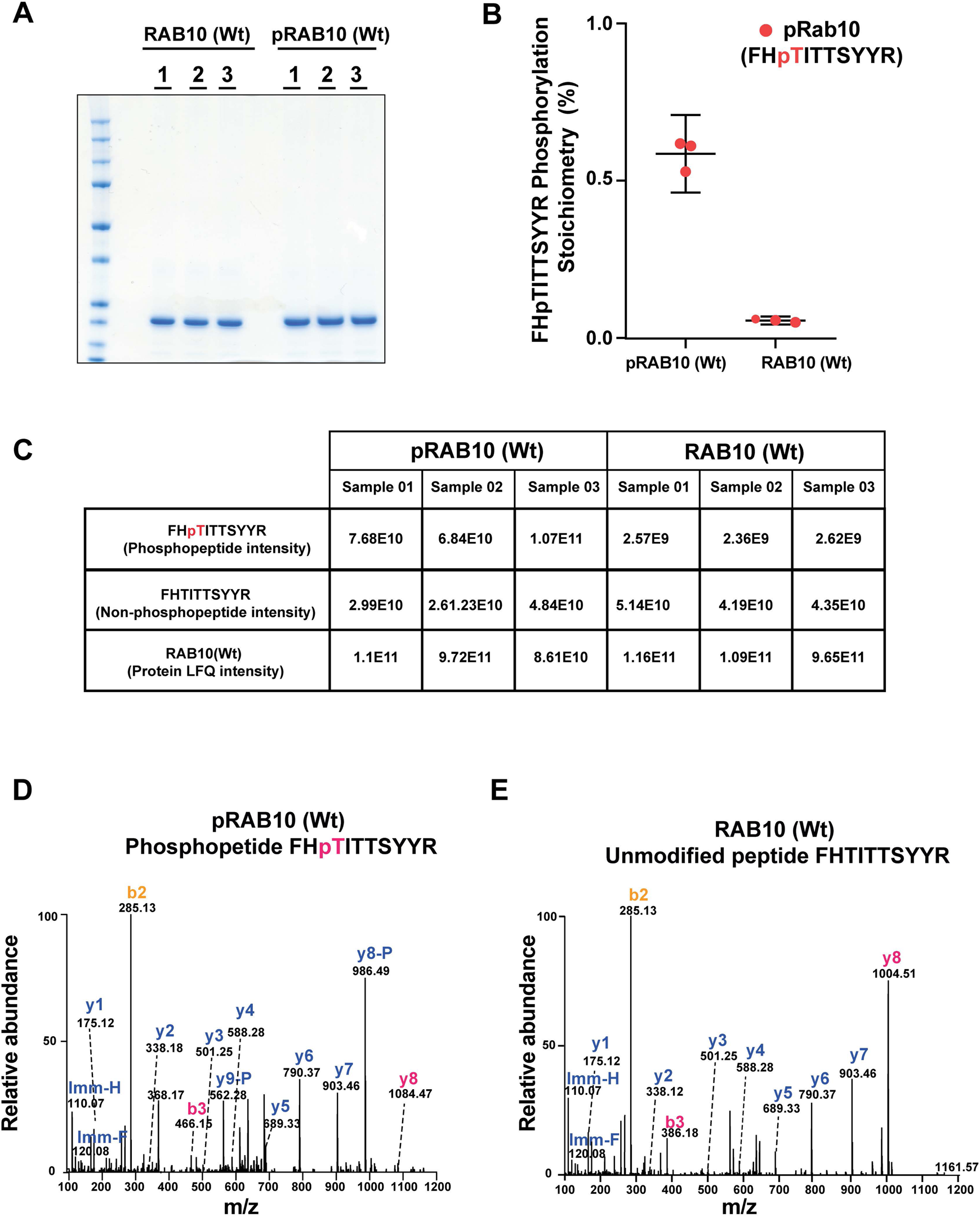
Mass spectrometry characterization of Thr73 phosphorylated Rab10[1-181, Wt]. (A) 2 µg of Rab10 [Wt] and pRab10 [Wt] in technical triplicates was loaded on a 4-12% Bis-Tris gradient gel. Following colloidal Coomassie staining, the lane encompassing 25 kDa region was excised and subjected to In-gel digestion using trypsin and analyzed on QE – HFX mass spectrometer. (B) and (C). The mass spectrometry raw data from Rab10 [Wt] and pRab10 [Wt] samples were searched using MaxQuant pipeline. The Phosphopeptide FHpTITTSYYR, unmodified peptide FHTITTSYYR intensities and Rab10 [Wt] and pRab10 [Wt] protein LFQ intensities as shown in table (C) were used to determine the FHpTITTSYYR, phosphorylation stoichiometry. To enable accurate phosphosite stoichiometry to be calculated both the modified and unmodified Thr73 peptides were excluded in determining the protein LFQ intensities. The phosphorylation stoichiometry in pRab10 [Wt] was found to be ∼60% and in Rab10 [Wt] was found to be <5% which is shown as separated scatter plot (B), the error bars represent the mean with 95% CI. (D) Representative annotated MS/MS spectrum of a Thr73, FHpTITTSYYR (m/z 684.80389; Z=2) identified in pRAB10 [Wt]. The m/z 466.15, z=1 of a b3 ion and m/z 1084.47, z=1 of y8 ions indicated with dotted lines confirms the phosphosite localization of the Phosphopeptide FHpTITTSYYR. (E) Representative annotated MS/MS spectrum of an unmodified peptide, FHTITTSYYR (m/z 644.82023; z=2) identified in RAB10 [Wt].

**Figure 7-Figure Supplement 6.**
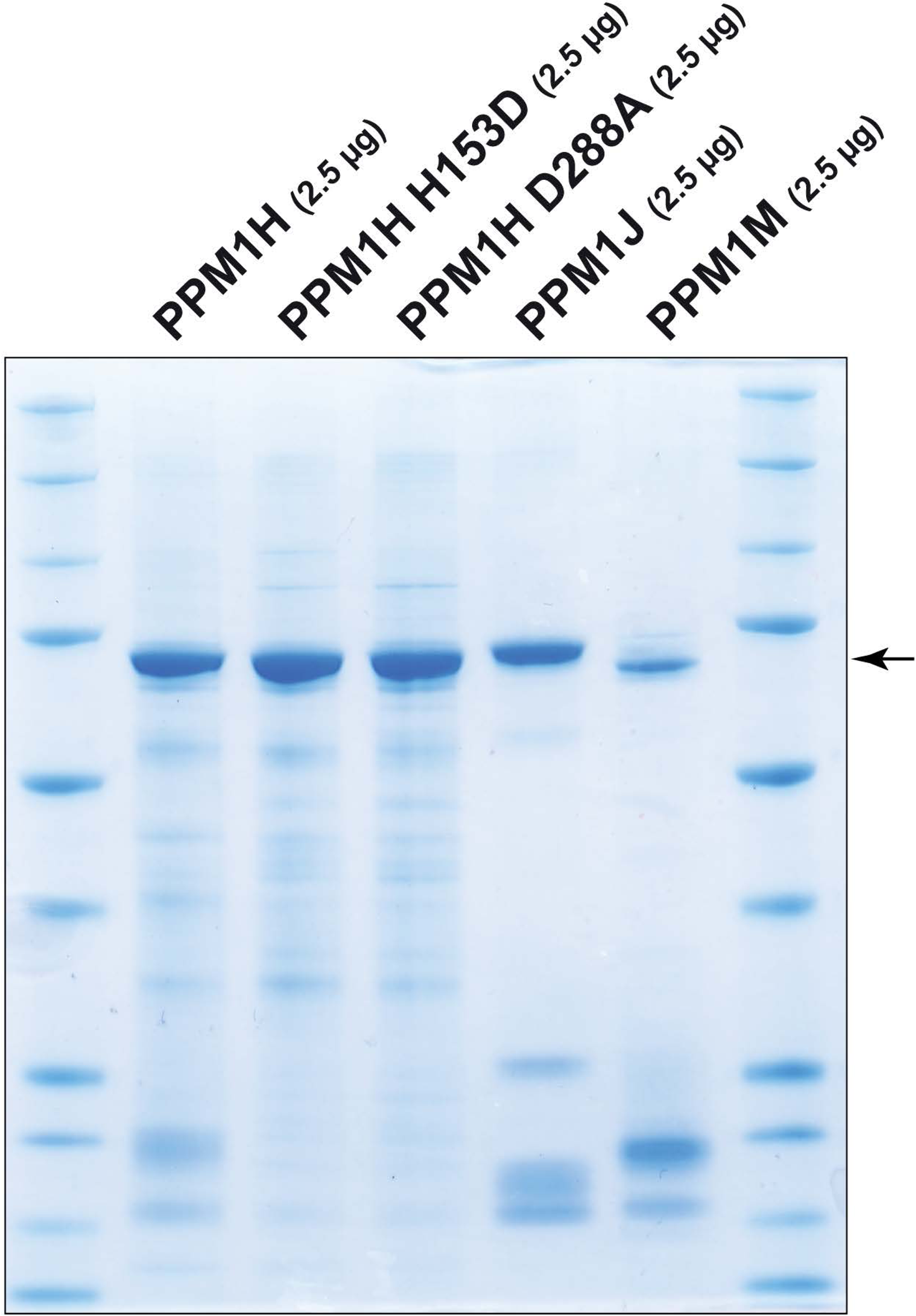
Coomassie stained 4-20% gradient SDS polyacrylamide gel electrophoresis analysis of 2.5 µg of purified recombinant phosphatases PPM1H, PPM1H H153D, PPM1H D288A, PPM1J and PPM1M (all with His-Sumo N-terminal tag, expressed in E. coli)

**Figure 8-Figure Supplement 1.**
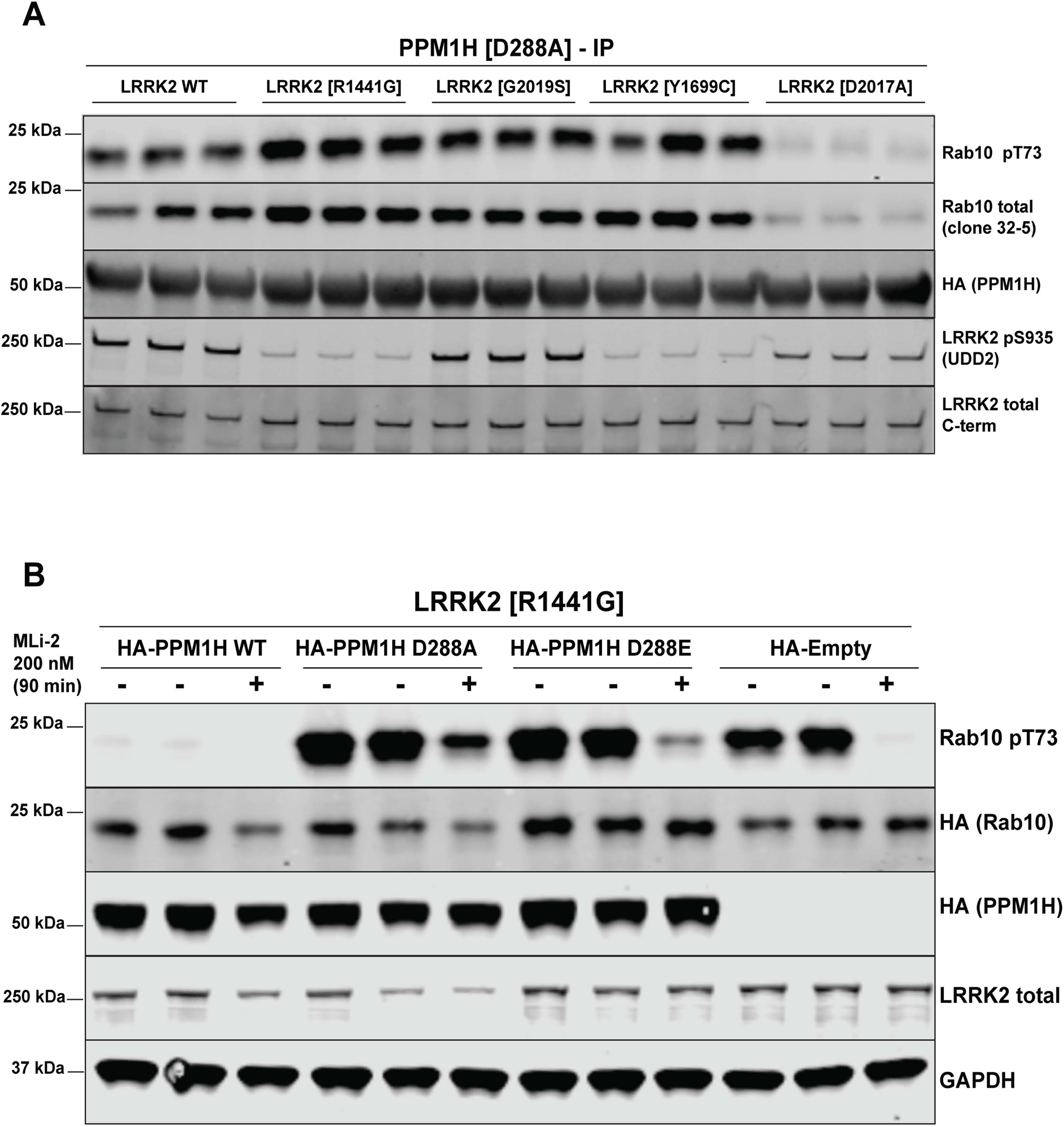
PPM1H[D288A] substrate trapping mutant stably interacts with LRRK2 phosphorylated Rab10 protecting it from dephosphorylation. (A) HEK293 cells were transiently transfected with constructs expressing PPM1H[D288A] with either wild type or indicated pathogenic or catalytically inactive LRRK2. 24 h post-transfection, cells were lysed and analyzed by immunoblotting with the indicated antibodies (1 µg/ml). Membranes were developed using Odyssey CLx Western Blot imaging. Each lane represents biological replicates. (B) As in (A) except that HEK293 cells were transiently transfected with constructs expressing Flag-LRRK2[R1441G] and either wild type HA-PPM1H or indicated HA-PPM1H mutants or empty HA-vector for no phosphatase control. 24h post-transfection cells were treated ± 200 nM MLi-2 prior to cell lysis for 90 min.

**Figure 8-Figure Supplement 2.**
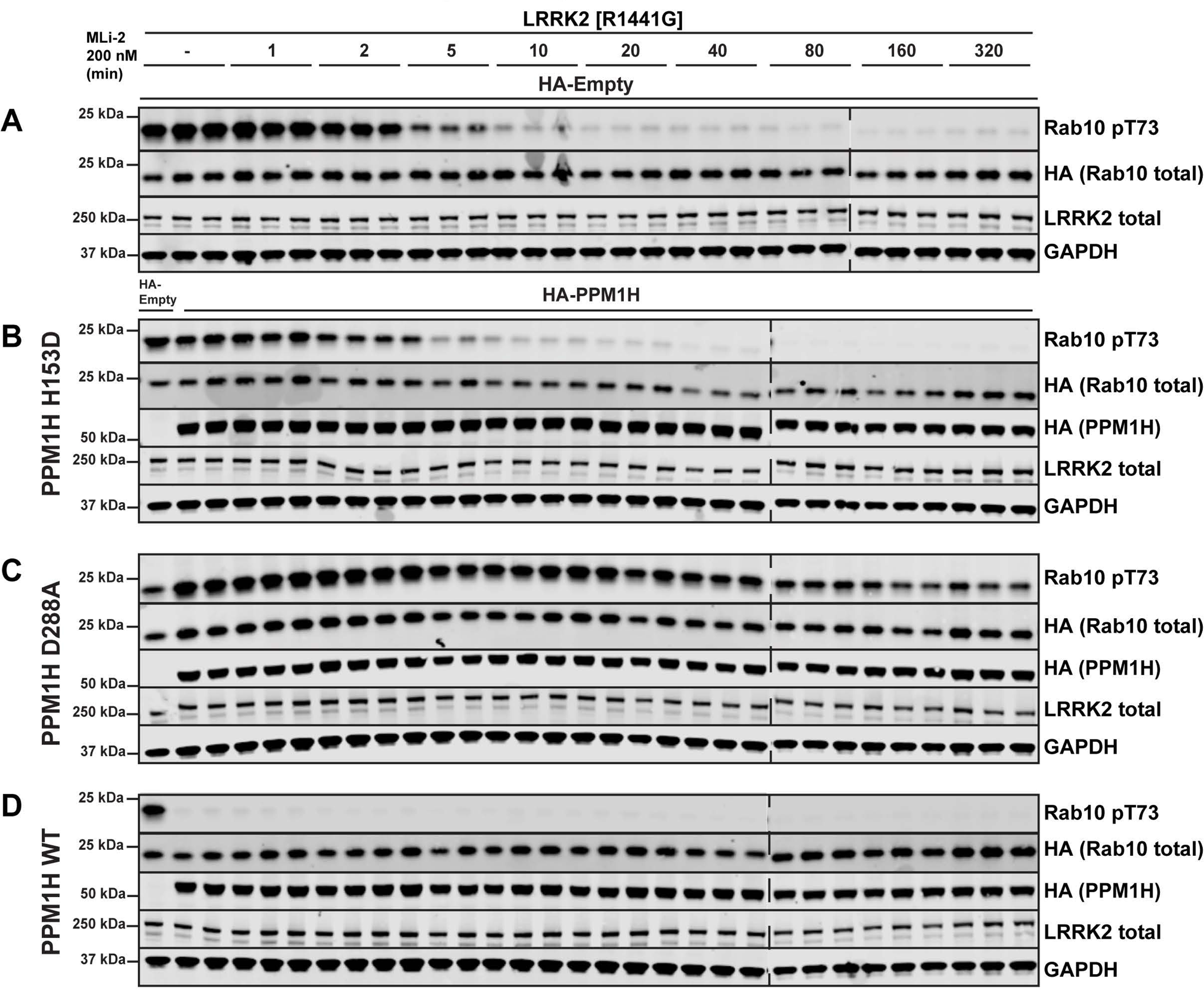
PPM1H[D288A] substrate trapping mutant blocks dephosphorylation of endogenous Rab10. HEK293 cells were transiently transfected with LRRK2[R1441G] and constructs expressing empty HA-vector (no phosphatase control) (A), catalytically inactive PPM1H[H153D] (B), substrate trapping PPM1H[D288A] (C) or wild type PPM1H (D). 24 h post-transfection, cells were treated with ± 200 nM MLi-2 for the time-points indicated, lysed and analyzed by immunoblotting with the indicated antibodies (1 µg/ml). Membranes were developed using Odyssey CLx Western Blot imaging. Each lane represents biological replicates.

**Figure 11-Figure Supplement 1.**
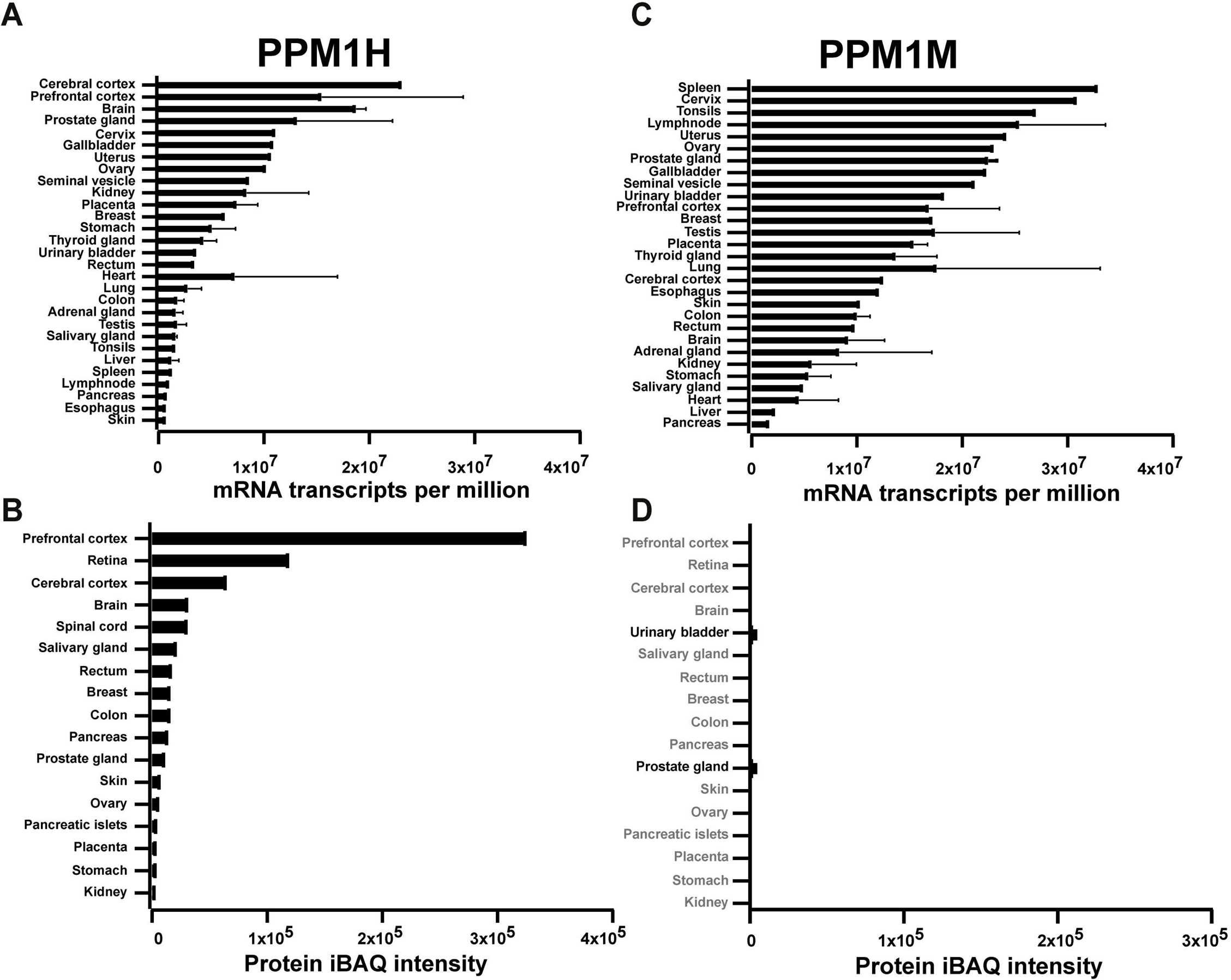
The abundance of PPM1H and PPM1M mRNA and protein in human tissues was analyzed using data downloaded from ProtemicsdB public repository database (https://www.proteomicsdb.org/) [67]. The median mRNA expression across the tissues studied is depicted as transcripts per million. The median protein expression for both PPM1H and PPM1M is calculated as the average normalized iBAQ (intensity based absolute quantification).

**Figure 11-Figure Supplement 2.**
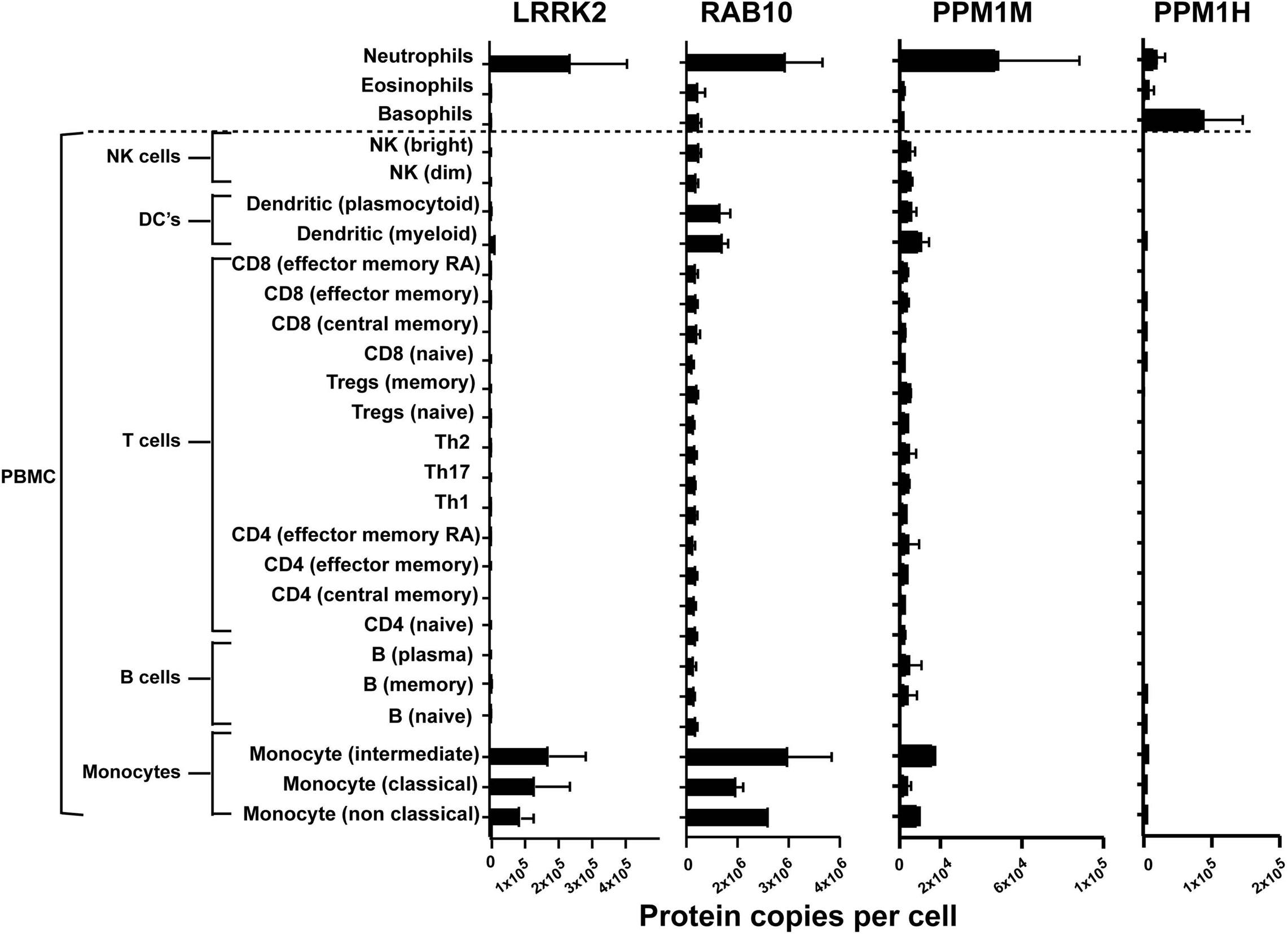
**The abundance of PPM1H and PPM1M proteins in immune cells isolated from human blood using data available from the immprot database (http://www.immprot.org) [68]**. The study utilized pure populations of immune cells isolated from human blood using fluorescence-activated cell sorting, and whole cell proteomics data were generated. Data were analyzed using the histone ruler to estimate protein copy numbers per cell. The graphs show the number of protein copies per cell for PPM1H, PPM1M, LRRK2 and RAB10 in a range of peripheral blood immune cells including subsets of T cells, B cells, monocytes, NK cells, dendritic cells and the granulocytes neutrophils, basophils and eosinophils.

